# The *C. elegans* endonuclease NUC-1 acts in engulfing cells to degrade the apoptotic cell DNA

**DOI:** 10.64898/2026.08.19.745295

**Authors:** Jonathan C. Pickett, Xianghua Liu, Lucia Chiao, Omar Cruz Ramirez, Lathan Lucas, Zheng Zhou

## Abstract

During *C. elegans* embryonic development, cells undergoing programmed cell death are engulfed by neighboring cells and degraded inside phagosomes. Here we characterize a DNase responsible for the degradation of the chromatin DNA of apoptotic cells. In the past, NUC-1, a homolog of mammalian DNase II, which is only active at acidic pH, was claimed to act in apoptotic cell nuclei for chromatin DNA degradation by some researchers, yet proposed to act in engulfing cells by others. We found that NUC-1 acts exclusively in engulfing cells to degrade apoptotic cell DNA. In *nuc-1* mutant embryos, apoptotic cell chromatin DNA remains undegraded. We observed that being engulfed is necessary for the apoptotic chromatin DNA to be degraded. In addition, specific expression of *nuc-1* in the engulfing but not dying cells rescues the *nuc-1* mutant phenotype. Furthermore, blocking the fusion between lysosomes and a phagosome in engulfing cells blocks apoptotic chromatin DNA degradation. NUC-1 was reported to be a lysosome-located enzyme. We not only confirmed this localization pattern, but also further determined that NUC-1 does not reside in the nuclei of either apoptotic or live cells. This, together with our finding that the nucleus of an apoptotic cell is not acidic, indicates that NUC-1 does not act in the apoptotic cell nucleus; rather, it acts in the engulfing cell phagosomal lumen to degrade apoptotic chromatin DNA. Our work clarified a long-standing controversy regarding the action of NUC-1 and advanced our knowledge of the mechanisms that drive the degradation of specific components of dying cells.

**Graphical Abstract:** 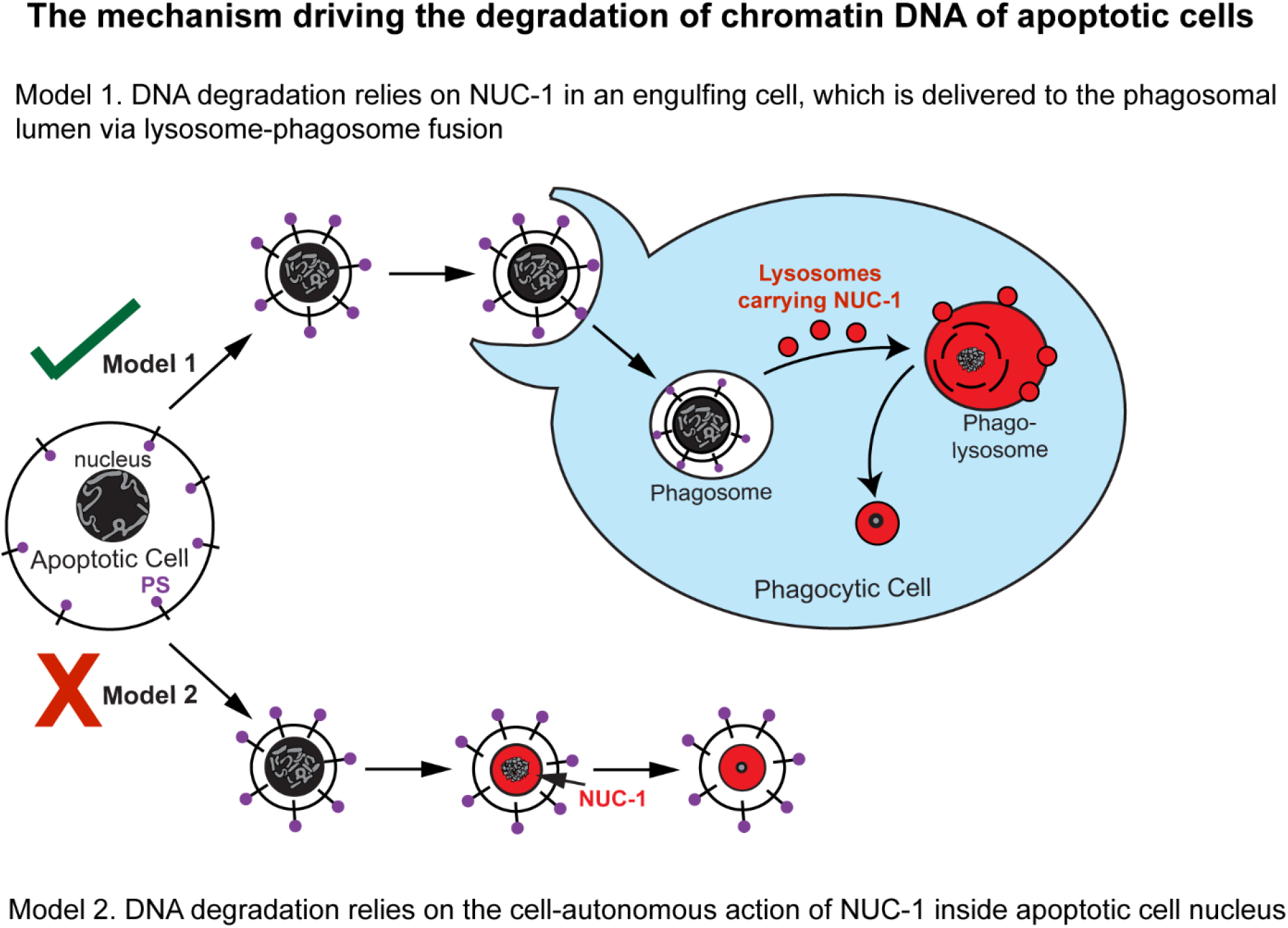

**Article Summary:** The degradation of apoptotic cell chromatin DNA relies on the sequential action of multiple DNases. *C. elegans* NUC-1, a mammalian DNase II homolog, is essential for apoptotic DNA degradation. Previously, there was a long-standing debate about whether NUC-1 acts in the apoptotic cells or their engulfing cells to complete DNA degradation. The authors report that NUC-1 does not act in the apoptotic cell nucleus; rather, it acts in the engulfing cell phagosomes to degrade apoptotic chromatin DNA. This work clarifies the controversy regarding NUC-1 action and sheds light on the conserved mechanisms that drive the degradation of dying cells’ components.

## Introduction

During the development of the soil nematode *Caenorhabditis elegans*, cells undergoing programmed apoptotic cell death are recognized by engulfing cells and subsequently engulfed into phagosomes, vacuoles sealed by membranes originating from the plasma membrane of the engulfing cells (Lu and Zhou 2012). Inside the phagosomal lumen, the lipids, proteins, carbohydrates, and nucleic acids of an apoptotic cell are degraded by hydrolases brought over by lysosomes that fuse to the phagosome (Lu and Zhou 2012; Levin, Grinstein, and Canton 2016). The fate of being engulfed and degraded is conserved for apoptotic cells in metazoans. In mice, defects in the degradation of chromatin DNA in apoptotic cells induce autoimmune responses that trigger chronic autoimmune diseases such as polyarthritis (Kawane et al. 2006). Knocking out DNase II, a lysosomal endonuclease, blocks the degradation of chromatin DNA expelled from erythroid precursor cells and apoptotic cells, resulting in embryonic lethality (Kawane et al. 2001; Kawane et al. 2003). Deciphering the mechanisms driving the degradation of apoptotic cell materials thus will improve our understanding of the development of autoimmune diseases and other disorders.

A loss-of-function mutation of *nuc-1*, which encodes the *C. elegans* homolog of DNase II, was identified by its lack of apoptotic cell DNA degradation (Sulston 1976). *In nuc-1(e1392)* loss-of-function mutant worms, programmed cell death and engulfment events are normal, yet the DNA of apoptotic cells in the posterior ventral cord of L1/L2 larvae remains as a pyknotic spot when worms were stained with Feulgen, a DNA dye (Sulston 1976; Hedgecock, Sulston, and Thomson 1983). This observation was later confirmed by Syto 11 staining of the *C. elegans* larvae (Lai et al. 2009). Robust endonuclease activity of partially purified NUC-1 from *C. elegans* protein extracts has also been detected (Hevelone and Hartman 1988), indicating that NUC-1 is a *C. elegans* DNase II ortholog. Like mammalian DNase II, NUC-1 is only active in acidic environments, between pH 4-5 (Hevelone and Hartman 1988). Hedgecock, Sulston, and Thomson (1983) further observed that in *ced-1* and *ced-2* mutants, which are defective in cell-corpse engulfment, apoptotic cells in the posterior ventral cord also remain undegraded. They thus proposed that NUC-1 acted inside the engulfing cell to digest the DNA of engulfed apoptotic cells.

Among a few distinct early features of apoptosis is the autonomous cleavage of apoptotic chromatin DNA into 180 bp fragments by the caspase-activated DNase (CAD) enzyme, an endonuclease that produces 3’-hydroxyl ends (Nagata et al. 2003), which can be detected by the Terminal deoxynucleotidyl transferase dUTP nick end labeling (TUNEL) technique (Gavrieli, Sherman, and Ben-Sasson 1992). Although the *C. elegans* homolog of mammalian CAD has not been identified, TUNEL-positive puncta were observed in wild-type embryos in a manner dependent on *ced-3*, the *C. elegans* caspase 3 (Wu, Stanfield, and Horvitz 2000). Furthermore, in *nuc-1* mutant embryos, the number of TUNEL-positive puncta is increased by approximately 30-fold, suggesting that NUC-1 acts to further degrade the fragments generated by the *C. elegans* CAD equivalent in apoptotic cells (Wu, Stanfield, and Horvitz 2000). This conclusion is consistent with the previous report (Sulston 1976) that NUC-1 is required for the degradation of apoptotic cell DNA.

Regarding whether NUC-1 acts in the apoptotic cells or engulfing cells to degrade apoptotic chromatin DNA, two sets of publications propose opposite models. As described above, by staining chromatin DNA with Feulgen, Hedgecock, Sulston, and Thomson (1983) found that engulfment is required for apoptotic cell DNA to be degraded. In support of this model, NUC-1 is observed to localize to lysosomes (Guo et al. 2010). On the other hand, by monitoring the number of TUNEL-positive puncta, Wu, Stanfield, and Horvitz (2000) observed that both the *nuc-1*-independent and the *nuc-1*-dependent TUNEL-positive puncta are the same in number in the *ced-2*, *ced-5*, *ced-6*, and *ced-10* mutant embryos as in wild-type embryos. Based on these observations, Wu, Stanfield, and Horvitz (2000) concluded that both the initial fragmentation of apoptotic chromatin DNA and the degradation of these fragments occurred cell autonomously. Later, Lai et al. (2009) and Yu et al. (2015) proposed that NUC-1 might be expressed in certain embryonic cells, secreted, internalized by apoptotic cells, and in this manner facilitate the degradation of apoptotic chromatin DNA.

The contradictory conclusions regarding whether NUC-1 acts in the engulfing or apoptotic cells to degrade apoptotic chromatin DNA are puzzling. We suspect that they might be results of DNA degradation detection methods and the different developmental stages of the *C. elegans* monitored. The TUNEL stain and the Feulgen stain require fixed specimens and are thus not suitable for monitoring the dynamics of apoptotic DNA degradation over time. In addition, the TUNEL staining positive signal only represents one particular state of DNA fragmentation.

Moreover, multiple reports (Wu, Stanfield, and Horvitz 2000; Lai et al. 2009; Yu et al. 2015) rely on a questionable assumption that in the *ced-1*, *ced-2*, *ced-5*, *ced-6*, *ced-7*, and *ced-10* mutant embryos, all apoptotic cells are engulfed to generate their conclusions. To clarify whether NUC-1 acts in an apoptotic cell or its engulfing cell to degrade apoptotic chromatin DNA fragments, we developed an *in vivo* fluorescent reporter for chromatin DNA. This reporter was used to monitor the dynamic chromatin DNA condensation and DNA degradation in time-lapse recording procedures during apoptosis in developing embryos. Our results indicate that engulfment is needed for NUC-1 to degrade apoptotic chromatin DNA. Furthermore, the fusion between lysosomes and the phagosome in an engulfing cell is essential for DNA degradation. Meanwhile, NUC-1 is localized to lysosomes but is not observed in the nucleus of an apoptotic cell. Together, our results indicate that NUC-1 acts in engulfing cells to execute its DNA-degradation activity within phagosomes.

## Materials and Methods

### Mutations, strains, and transgenic arrays

*C. elegans* strains were grown at 20°C as previously described (Riddle et al. 1997) unless indicated otherwise. The N2 Bristol strain was used as the wild-type control strain. Mutations are described in Riddle et al. (1997) and by Wormbase (http://www.wormbase.org) unless noted otherwise (**Table S1**): LG1: *ced-1(e1735)*; LGII: *rab-7(ok511)*; *ced-5(n1812)*; LGV: *unc-76(e911)*; LGX: *nuc-1(e1392).* The *rab-7(ok511)* homozygous strains are maternal-effect embryonic lethal, and the mIn1 balancer maintained this allele with an integrated pharyngeal GFP marker (**Table S1**) (Edgley et al. 2006). To obtain *rab-7(ok511)* m^-^z^-^ homozygous embryos, GFP^-^ *rab-7(ok511)* homozygous hermaphrodites were isolated among the progeny of the strain VC308, and their progeny were collected as embryos.

Extrachromosomal arrays were generated by the microinjection of plasmids with the co-injection marker p76-18B [*punc-76(+)*] into *unc-76(e911)* mutants (Bloom and Horvitz 1997; Jin 1999). Non-Unc mutants were identified as transgenic animals. Integrated transgenic arrays were generated by gamma irradiation (Jin 1999). Integrated arrays generated in this study are as follows (**Table S1**): LGI: *enIs71[Pced-1PH(hPLCγ)::mrfp* and P*his-72 his-72::mCherry]*.

### Plasmid construction

P*ced-1 nuc-1::gfp* (pXL57) and P*ced-1 nuc-1::mCherry* (pXL63) were reported in (Haley, Wang, and Zhou 2018). P95.75_mNeonGreen (pOPR15), P*his-72 his-72::mCherry* (pXL75), P*his-72 his-72::gfp::mCherry* (pXL68), and P*ced-1 mCherry::lgg-1* (pZZ1091) were reported in (Peña-Ramos et al. 2022). *Pced-1 ctns-1::gfp* (pBZ212) was reported in (Yu, Lu, and Zhou 2008). To generate P*efn-2 nuc-1::gfp* (pOCR15), the 2.8-kb piece of DNA 5’ of the start codon was PCR amplified from N2 worm extract, using the primer pair OC79/OC80 (**Table S1**). This fragment was used to replace P*ced-1* in P*ced-1 nuc-1::gfp*. To generate P*efn-2 his-72::mCherry* (pJP14), the *his-72::mCherry* cassette was obtained from *Phis-72 his-72::mCherry* by digestion with BamH1 and Apa1 and replaced the *nuc-1::gfp* cassette in P*efn-2 nuc-1::gfp*. *Phis-72 nuc-1::mCherry* (pJP3) was constructed by PCR amplification of *Phis-72* from P*his-72 his-72::gfp::mCherry* using the primer pair ZZ1227/ZZ1228 followed by digestion with Sph1/BamH1 and replacing P*ced-1* in P*ced-1 nuc-1::mCherry*. To generate P*his-72 ctns-1::gfp* (pJP4), P*his-72* was PCR amplified from P*his-72 his-72::mCherry* using the primer pair ZZ1268/ZZ1269, digested with Sph1 and Sal1, and inserted into P*ced-1 ctns-1::gfp*, replacing P*ced-1*. P*nuc-1* was PCR amplified from N2 worm extract using the primer pair ZZ1036/ZZ1037 and digested with Sph1 and BamH1. It was then cloned into pOPR15, a vector that carries mNeonGreen (mNG), to generate P*nuc-1 mNG.* P*nuc-1 nuc-1::mNG* (pJP1) was generated by inserting an Age1-Xma1 fragment of full-length *nuc-1* cDNA into P*nuc-1 mNG.* P*nuc-1 nuc-1::mStayGold* (pJP17) was generated by replacing mNG with an Age1-Apa1 fragment containing mStayGold, a monomeric form of StayGold with *C. elegans* codon optimization, contained in pSM-mStayGold (a gift from Kota Mizumoto (Addgene plasmid # 234317)) (Ko and Mizumoto 2025). To construct *Pced-1 mCherry::nuc-1* (pZZ1251), *nuc-1* cDNA was PCR-amplified from P*ced-1 nuc-1::gfp* using the primer pair ZZ1330/ZZ1331, digested with Xma1 and Kpn1, and cloned into the Xma1 and Kpn1 sites of P*ced-1 mCherry::lgg-1* (Peña-Ramos et al. 2022), replacing the *lgg-1* cDNA. *Pced-1 mCherry::(Δss)nuc-1* (pZZ1254) was constructed by PCR amplification of *nuc-1* cDNA using the primer pair ZZ1332/ZZ1331, starting from the 64 bp after ATG, followed by digestion with Xma1/Kpn1 and cloning into the Xma1 and Kpn1 sites of P*ced-1 mCherry::lgg-1*, replacing the *lgg-1* cDNA.

### Fluorescence microscopy and time-lapse imaging

A DeltaVision Elite Deconvolution Imaging System (GE Healthcare, Inc.) equipped with a DIC imaging apparatus and a Photometrics Coolsnap 2 digital camera was used to capture fluorescence, and DIC images Applied Precision SoftWoRx 5.6 software was utilized for deconvolving and analyzing the images (Lu et al. 2009). During the clearance process of cell corpses C1, C2, and C3, embryos were monitored on their ventral surface starting at ∼310 min post-first cleavage using an established time-lapse imaging protocol (Lu et al. 2009). To observe the clearance of Cm, C4, and C5, the ventral surface of an embryo was monitored starting at ∼240 min post-first cleavage. For all the time-lapse recording programs, 12 to 16 serial *Z-* sections (at 0.5-μm intervals) were captured every 2 or 3 min, with recordings typically lasting between 60 and 180 min.

#### To determine the time point when apoptotic cell engulfment is complete

We used a few different methods to determine the time point when apoptotic cell engulfment is complete, the “0 min” time point, depending on the marker available. When available, PH(PLCγ)::GFP (Shen et al. 2013) or CED-1::GFP (Lu et al. 2009), expressed in engulfing cells under P*ced-1*, are pseudopod reporters that are monitored to determine the time point when the pseudopods are sealed, representing the “0 min” time point. When the fluorescence channels are reserved for other reporters, the first moment the button-like appearance arises in the DIC channel is considered the “0 min” moment, as our experiments (**Fig S1**) have shown that the rise of the DIC cell corpse appearance coincides with the “0 min” time point.

#### To measure the degradation of chromatin DNA of apoptotic cells Cm, C3, C4, and C5

We measured the size of the nucleus of engulfed Cm, C3, C4, or C5 over time by following the ubiquitously expressed P*his-72 his-72::mCherry* reporter as described in (Peña-Ramos et al. 2022). Briefly, the diameter at each time point (Dt) of the HIS-72::mCherry^+^ patch inside a phagosome was measured as a representation of chromatin DNA. The DNA degradation index (DDI) is calculated as the ratio Dt/D0. Once DDI was reduced to < 0.5, the chromatin DNA is considered degraded.

#### To measure the efficiency of chromatin DNA degradation inside phagosomes in 1.5-fold stage embryos

HIS-72::GFP::mCherry was used in this assay for distinguishing engulfed apoptotic cell nuclei from unengulfed apoptotic cell or live cell nuclei. Only when deposited into the phagosomal lumen for at least 20 min would the GFP signal significantly reduce due to quenching of the GFP fluorophore by the acidic pH. The GFP and mCherry signals from wild-type or *nuc-1* mutant embryos at 1.5-fold stage were recorded in serial z-sections, with 40 z slices at the 0.5 μm z interval. The numbers of mCherry^+^ GFP^-^ patches with diameters >1.0 μm, which represent chromatin DNA of engulfed apoptotic cells that were not efficiently degraded, were counted throughout all 40 z slices.

In *rab-7(ok511)* mutant embryos, the acidification of the phagosomal lumen is defective (Fig. S4 C-F). As a consequence, HIS-72::GFP signal does not diminish inside the lumen. We thus relied on the condensed morphology of HIS-72::mCherry and the button-like morphology to distinguish apoptotic cells inside phagosomal lumen. With this modification, the samples we monitored include nascent phagosomes, not those phagosomes that were at least 20-min old.

#### To score the recruitment and entry of NUC-1::mCherry into the phagosomal lumen

In this assay, NUC-1::mCherry was exclusively produced in engulfing cells under the control of P*ced-1*. The surfaces of the nascent phagosomes are labeled by co-expressed PH(PLCγ)::GFP. The time point when the GFP-labeled pseudopods seal as a ring is counted as the “0 min” time point, when engulfment is just complete. The first moment when an mCherry puncta is attached to the phagosomal surface is counted as the time point when NUC-1 recruitment starts. The first moment when the mCherry is obvious in the middle of phagosomal lumen is counted as the time point when NUC-1 entry starts.

#### To quantify the acidification status of the phagosomal lumen or cell nucleus

This assay uses the HIS-72::GFP::mCherry fusion protein expressed in apoptotic cells under the control of P*his-72* as the acidification reporter, taking advantage of the low-pH-sensitive and resistant nature of GFP (*pka* = 6.0) and mCherry (*pka* = 4.5) (Peña-Ramos et al. 2022; Peña-Ramos and Zhou 2023). At each time point, the mCherry and GFP signal intensities of a fixed area (3×3 pixels) in the center of a phagosome or a nucleus (IntmCherry, IntGFP) were recorded.

The acidification status (AS) at a particular time point is defined as IntGFP / IntmCherry. The relative acidification index (RAITn) at a particular time point in relation to the “0 min” time point is defined as ASTn / AST0. The RAITn value 1.0 indicates no acidification of the phagosome compared to the “0 min” value.

## Results

### Apoptotic cell DNA remains undegraded in *nuc-1* mutants

Histone H3 is part of the central core of nucleosomes, the primary structural units of chromatin (McGinty and Tan 2015). To monitor the status of chromatin DNA in apoptotic cells, we generated an mCherry-tagged histone H3 (*his-72*::mCherry) reporter expressed under the *his-72* promoter (P*his-72*) as a chromatin DNA reporter (Pena-Ramos et al, 2022). The HIS-72::mCherry reporter in a cell forms an mCherry^+^ patch (Fig. 1 C, E, G). We used time-lapse imaging to monitor the size of the mCherry^+^ patch in the apoptotic cells listed in Fig. 1(A, B) over time in wild-type and *nuc-1(e1392)* null mutant strains (**Materials and Methods**). In wild-type, we first monitored three dynamic events occurring when a cell underwent apoptosis: the condensation of chromatin DNA (via HIS-72::mCherry), the appearance of the button-like morphology (via DIC microscopy), and the engulfment by a neighboring cell (via engulfing cell-expressed PH(PLCγ) domain::GFP, a plasma membrane marker (Shen et al. 2013)) (Fig. S1) (**Materials and Methods**). In 87.5% of the apoptotic C3s monitored, both chromatin condensation and the appearance of the button-like DIC morphology occur within a 4-min time range, from 2 min before to 2 min after the completion of engulfment (the “0 min” time point) (Fig. S1). Because these three events occurred nearly simultaneously, in the strains where PH::GFP was unavailable to be used as a reporter for the engulfment status of an apoptotic cell, the appearance of the button-like DIC morphology was used as the “0 min” time point, except in engulfment-defective mutants.

**Figure 1.**
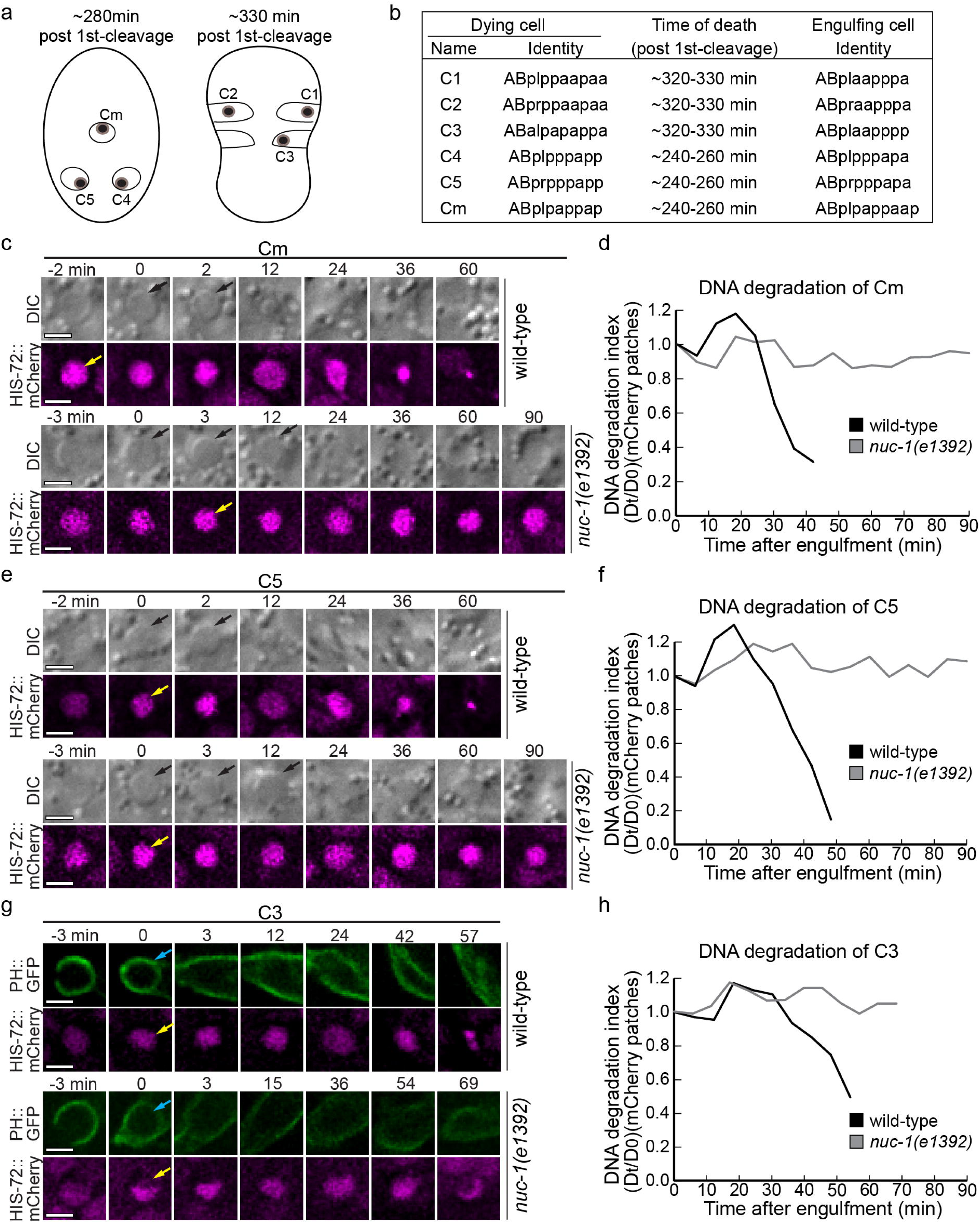
Time-lapse recording reveals that the degradation of apoptotic cell DNA is severely defective in *nuc-1* mutant embryos. (A) The positions of the dying (dots) and corresponding engulfing cells (open circles or petals) that we monitored with time-lapse recording on the ventral side, at the moment when engulfment occurs. Anterior is towards the top. (B) The identity of the dying cells and their engulfing cells monitored in this report. (C, E, G) Time-lapse images monitoring the chromatin DNA of Cm (C), C5 (E), and C3 (G) before and after their engulfment with HIS-72::mCherry in wild-type and *nuc-1* mutant embryos. The corresponding DIC images in C and E and the PH(PLCγ)::GFP images in (G) are for determining the moment that engulfment is complete, the “0 min” time point. Black arrows on DIC images mark the cell corpse. Yellow arrows on mCherry images mark chromatin DNA at the moment when it starts to appear condensed. Blue arrows in G mark the nascent phagosomes containing C3. Scale bars are 2 μm. (D, F, H) The diameters of the HIS-72::mCherry^+^ patches in Cm (C), C5 (E), or C3 (G) are measured over time and compared to that at “0 min”. The Dt/D0 ratio is considered the DNA degradation index and is plotted over time. **Figure 1 ALT TEXT:** Images, table and graphs related to NUC-1 activity, with subfigures labelled from a to h showing apoptotic cells in *C. elegans* embryos and the clearance pattern of cell-corpse DNA in wild-type and *nuc-1* null mutant backgrounds.

We then monitored the degradation of apoptotic chromatin DNA inside phagosomes by measuring the diameter of the mCherry^+^ patches over time (Fig. 1) (**Materials and Methods**). As examples in Fig. 1(C-H) show, in wild-type embryos, the diameters of the mCherry^+^ patches in Cm (C-D), C5 (E-F), and C3 (G-H) all shrink to <50% of the “0 min”-size within 60 min. Conversely, in *nuc-1(e1392)* mutant embryos, the diameters of Cm, C5, and C3 remain the same as the “0 min”-size by the 90-min recording time (Fig. 1(C-H)). Quantitative analysis of multiple samples in *nuc-1* mutants (Fig. S2) demonstrates that the chromatin DNA of these cells inside phagosomes persists instead of being degraded.

To further examine whether the defect in DNA degradation occurs in all embryonic apoptotic cells in *nuc-1* mutants, we established another DNA degradation assay based on a HIS-72::GFP::mCherry double-colored reporter (**Materials and Methods**). The fluorophore in GFP is sensitive to acidic pH (Tsien 1998) inside the phagosome lumen, whereas the mCherry fluorophore is relatively acid-resistant (Shaner et al. 2004). An apoptotic cell that is engulfed inside a phagosome (Fig. 2A, yellow arrow or white arrowhead), which is mCherry^+^ GFP^-^, can be distinguished from an mCherry^+^ GFP^+^ live cell (white arrows) (Fig. 2A). When we measured the diameter of the mCherry^+^ GFP^-^ patches in the 1.5-fold stage embryos, in wild-type embryos we mostly found tiny (diameter <0.5 μm) puncta, which represent remnants of the apoptotic chromatin (Fig. 2A yellow arrows, 2B). In contrast, in *nuc-1(e1392)* mutant embryos, on average 20 mCherry^+^ GFP^-^ patches with diameters > 1 μm (Fig. 2A white arrowheads, 2B) were detected, indicating that chromatin DNA of apoptotic cells inside phagosomes was not efficiently degraded and that this defect affects all apoptotic cells, not just the Cm, C3, and C5 cell corpses that we measured (Figs. 1 and S2). We named this assay “DNA degradation assay in whole embryos”. Together, these findings indicate that in *nuc-1* mutants, the chromatin DNA of apoptotic cells is not efficiently degraded despite being normally engulfed.

**Figure 2.**
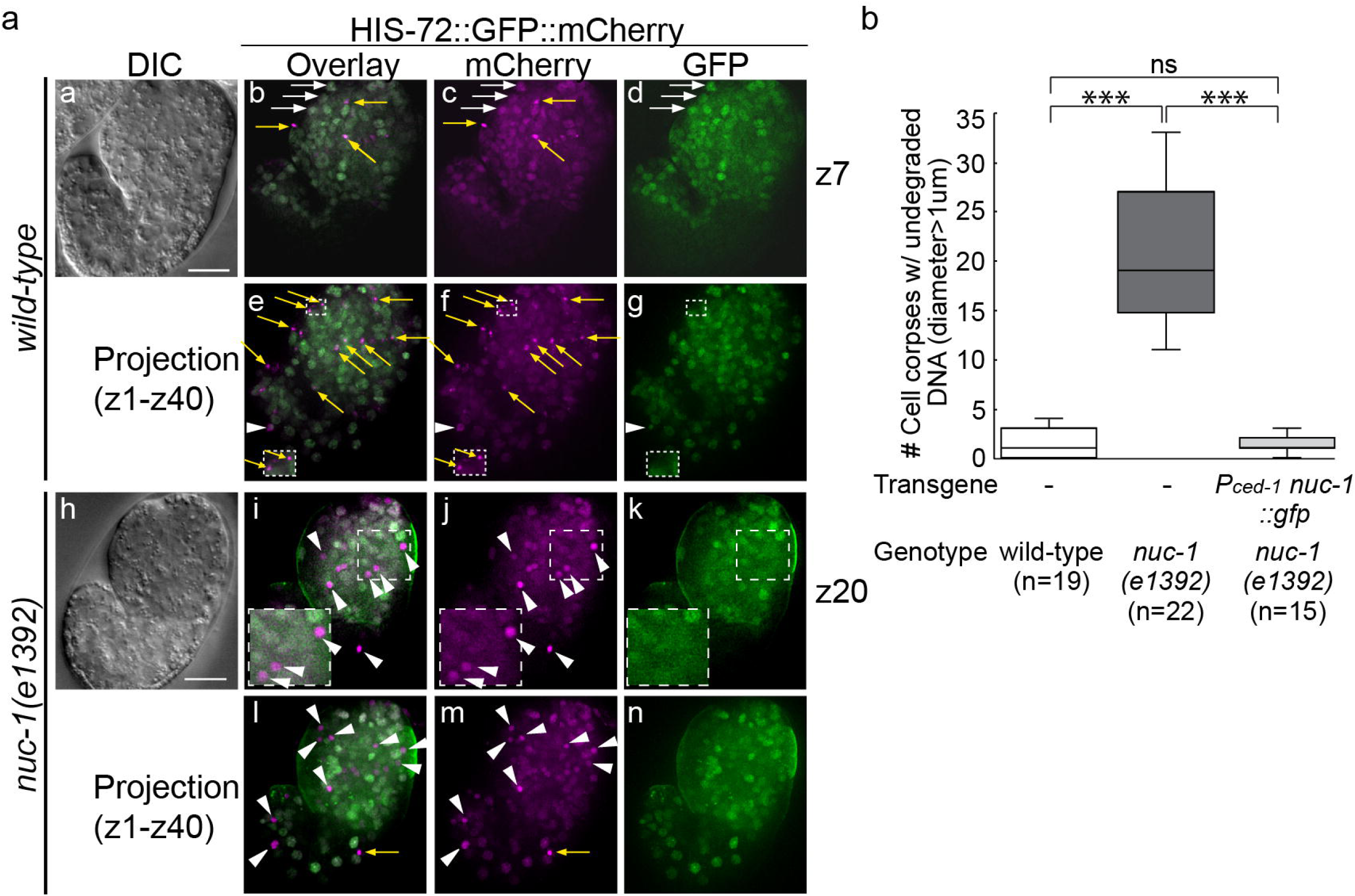
1.5-fold stage *nuc-1* mutant embryos carry many more cell corpses with undegraded chromatin DNA than wild-type embryos. (A) One each of wild-type (a-g) and *nuc-1(n1392)* mutant (h-n) embryos carrying a P*his-72* HIS-72::GFP::mCherry reporter. z, the position of an image in an optical section series of 40 slices with 0.5 μm interval. Nuclear DNA of live cells (white arrows) are mCherry^+^ and GFP^+^, and that of apoptotic cells inside mature phagolysosomes are mCherry^+^ and GFP^-^. Examples of those mCherry^+^ and GFP^-^ spots that are < 1μm in diameter are labeled with yellow arrows, and those >1μm in diameter labeled with white arrowheads. Regions framed with white boxes of dashed lines are the ones enlarged 1.5×1.5-fold in the inlets. Panels (e-g) and (l-n) display projected images of all 40 slices. Scale bars are 10μm. (B) Box-Whiskers plots of the number of apoptotic cells inside phagosomes with undegraded DNA (mCherry patch >1μm diameter) in 1.5-fold wild-type and *nuc-1(e1392)* embryos, and in *nuc-1(e1392)* embryos carrying the P*ced-1 nuc-1::gfp* transgene. n, number of embryos scored. “***”, “***”: p-value <0.001, Student *t*-test. **Figure 2 ALT TEXT:** Images and graph related to NUC-1 activity, with subfigures labelled a and b showing cells in 1.5-fold embryos to highlight different cell corpse clearance states in wild-type and *nuc-1* null mutant backgrounds.

### Engulfment is essential for the degradation of the chromatin DNA of an apoptotic cell

To determine whether NUC-1-dependent degradation of apoptotic cell DNA is required in dying or engulfing cells, we first examined whether engulfment is necessary for DNA degradation. In previous reports, apoptotic cells were assumed to remain unengulfed, that is, not internalized by any cell, in *ced-1*, *ced-2*, *ced-5*, *ced-6*, *ced-7*, and *ced-10* mutant embryos (Wu, Stanfield, and Horvitz 2000; Lai et al. 2009; Yu et al. 2015). However, we found this was incorrect, particularly in the mid-stage (1.5-fold) embryos. By monitoring the engulfment and degradation processes of a particular apoptotic cell in the wild-type, *ced-1*, and *ced-5* mutant strains, we previously found that although engulfment was slow and less efficient in the *ced-1* and *ced-5* mutants, a large number of cell corpses were eventually engulfed (Yu, Lu, and Zhou 2008). As a result, in mid-stage *ced-1* or *ced-5* mutant embryos, a substantial percentage of engulfed cell corpses are present among all cells that display the DIC-button-like morphology (Yu et al., 2008).

To investigate the status of engulfment of individual apoptotic cells more extensively, we further monitored, using time-lapse microscopy, the engulfment status of apoptotic cells Cm, C3, C4, and C5 in *ced-1(e1735)*, *ced-2(n1994)*, and *ced-5(n1812)* mutant embryos (Fig. S3) (**Materials and Methods**). We found that in all these mutants, engulfment was only partially defective, and substantial percentages of apoptotic cells were still engulfed (Fig. S3A). In addition, apoptotic cells of different identities were differentially sensitive to each mutation (Fig. S3A). Moreover, for those apoptotic cells that were eventually engulfed, the engulfment process was often significantly delayed (Fig. S3B). Thus, engulfed or unengulfed apoptotic cells with the same identity can be recognized and monitored over time from different embryos at the same stage. This partial engulfment defect allowed us to compare the efficiency of chromatin DNA degradation in engulfed versus unengulfed apoptotic cells.

In *ced-5(n1812)* null mutant embryos, we performed the time-lapse DNA degradation assay, first determining whether each apoptotic cell C4 and C5 was engulfed over time using a co-expressing PH::GFP reporter, while monitoring the size reduction of the HIS-72::mCherry^+^ patch. We found that the chromatin DNA within unengulfed C4 or C5 cell corpses never got degraded, even after the cell corpses were shed into the embryonic cavity (Fig. 3). Particularly, in 100% of the unengulfed C4 cell corpses, 90 min after engulfment, the diameters of the mCherry^+^ patches remain >90% of their original sizes, indicating a lack of DNA degradation (Fig. 3D). On the contrary, the diameters of the mCherry^+^ patches in 100% of the engulfed C4 cell corpses are reduced to 50% or less of their original diameters within 50 min after engulfment (Fig. 3D). These results strongly indicate that being engulfed is a pre-requisite for the apoptotic chromatin DNA to be degraded.

**Figure 3.**
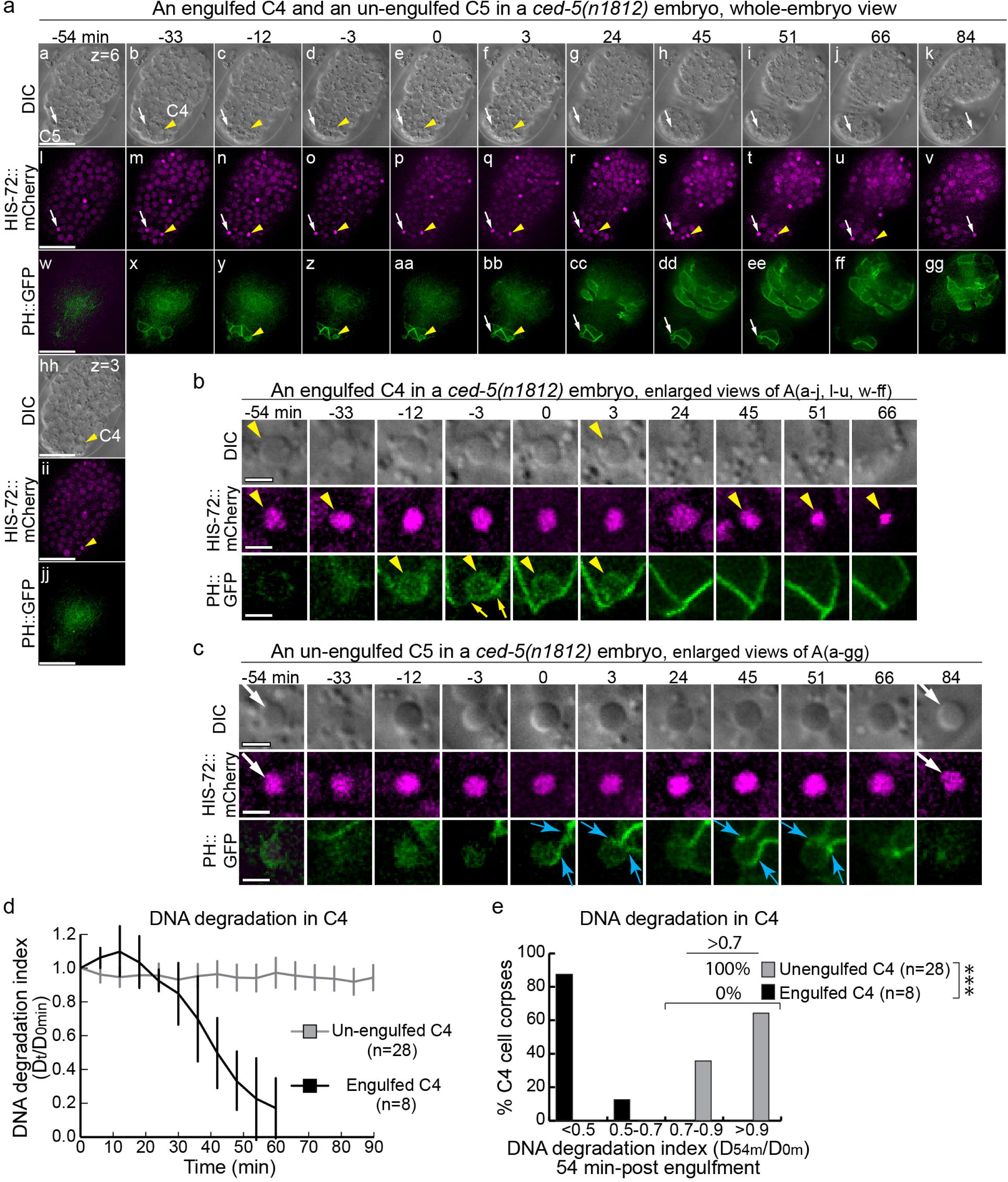
The degradation of apoptotic cell DNA depends on their engulfment. (A-C) Time-lapse recording images of the ventral side of a *ced-5(n1812)* mutant embryo, monitoring the engulfment of apoptotic cells C4 (yellow arrowheads) and C5 (white arrows) with PH(PLCγ)::GFP and the shrinkage of their chromatin DNA with HIS-72::mCherry. “0 min” is the moment the engulfment of C4 is completed. (A) The whole-embryo view. Scale bars are 15 μm. (B) Enlarged images of (A) around the engulfed C4 (The engulfment of this C4 is long delayed, judging by the embryonic developmental landmarks and the time point when the button-like DIC morphology appears). Yellow arrows label the two sides of the plasma membrane of the engulfing cell for C4. Scale bars are 2 μm. (C) Enlarged images of (A) around the unengulfed C5. Blue arrows label the pseudopods extending from the engulfing cell but later retracted. Scale bars are 2 μm. (D) The graph of the DNA degradation index of C4 over time, which is represented by the mean ratio of the diameter of the HIS-72::mCherry-labeled patch at each time point (Dt) after the completion of engulfment (“0 min”) over that of the “0 min” time point (D0). Error bars represent standard deviation (sd). n: the number of embryos analyzed. (E) A histogram of the DNA degradation index of C4 at 54 min-post “0 min” time point, based on the data presented in (D). “***”: p-value <0.001, Student *t*-test. **Figure 3 ALT TEXT:** Images and graphs related to cell corpse clearance in an engulfment mutant background, with subfigures labelled from a to e showing both engulfed and unengulfed cell corpses in the same engulfment mutant background.

### NUC-1 acts in engulfing cells to catalyze the degradation of apoptotic cell DNA

We tested whether expression of *nuc-1* under an engulfing-cell-specific or dying-cell-specific promoter would rescue the DNA degradation phenotype of a *nuc-1(e1392)* mutant. P*ced-1 nuc-1::gfp*, which is specifically expressed in engulfing cells under the control of the *ced-1* promoter (Zhou, Hartwieg, and Horvitz 2001b), completely rescues the apoptotic chromatin degradation defect in the “DNA-degradation-in-whole-embryo-assay”, when 1.5-fold stage embryos were scored for the presence of mCherry^+^ GFP^-^ chromatin patches with diameters >1.0 μm (Fig. 2B). Moreover, in the time-lapse DNA degradation assay, monitoring the reduction of the size of mCherry^+^ patches in Cm and C3, P*ced-1 nuc-1* also demonstrated efficient rescue of the *nuc-1(e1392)* phenotype (Fig. 4 A-D, G). On the other hand, when *nuc-1* is expressed under the control of P*enf-2*, a promoter that is specifically expressed in the mother of C3 (Grossman, Giurumescu, and Chisholm 2013) and C3 (Fig. 4H) but not in ABplaapppp, the engulfing cell for C3 (Fig. 4H), no significant rescue of the *nuc-1(e1392)* phenotype is observed (Fig. 4 E-G).

**Figure 4.**
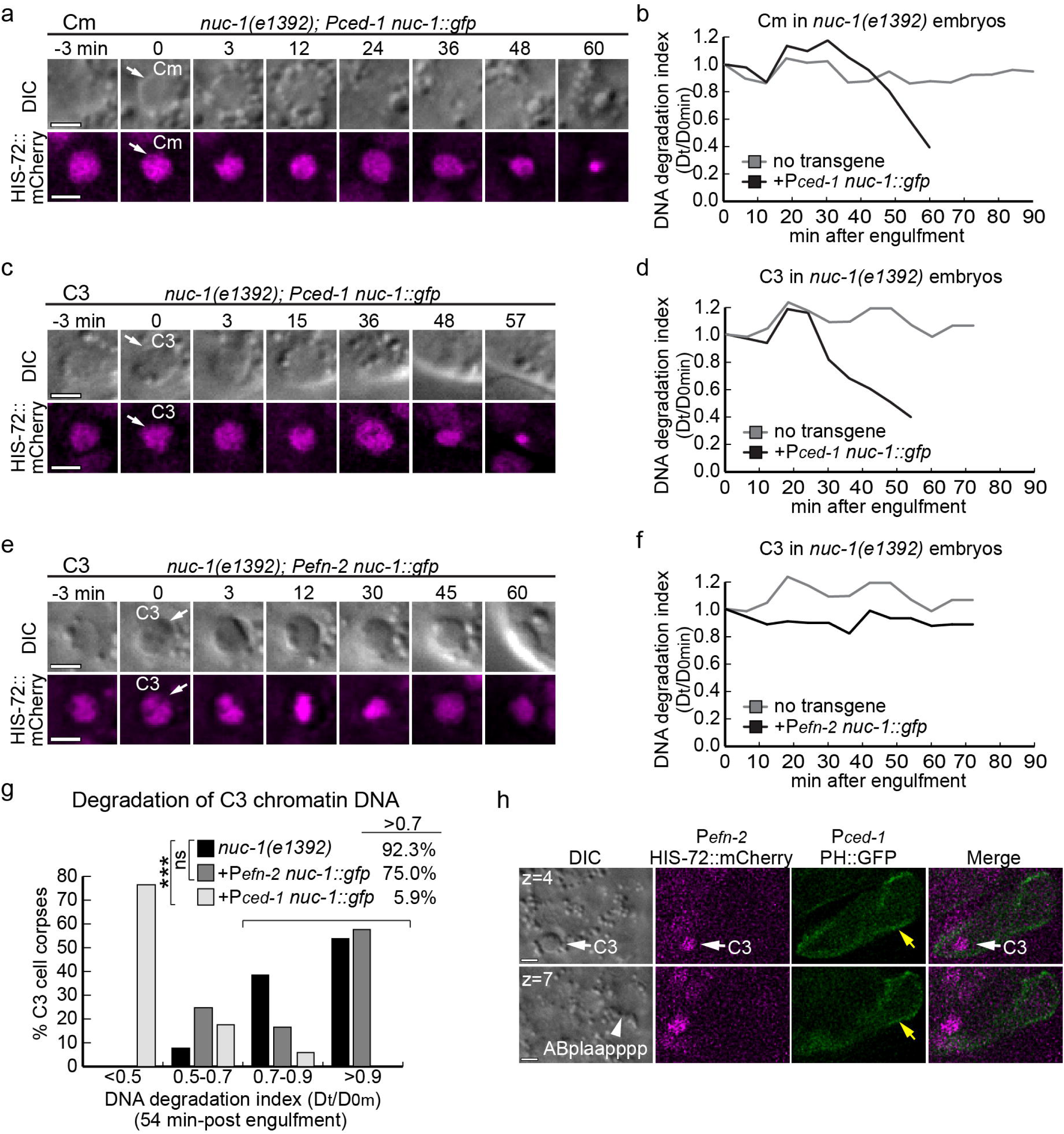
The expression of *nuc-1* in engulfing, but not in dying cells, fully rescues the apoptotic cell DNA degradation defect. (A, C, E) Time-lapse images monitoring the HIS-72::mCherry patches (white arrows) and the corresponding DIC images in Cm (A) and C3 (C and E) before and after the completion of engulfment (“0 min”), which is indicated by the 1^st^ appearance of the button-like DIC morphology (white arrows) in embryos expressing the *nuc-1*-rescuing constructs under P*ced-1* (A, C) and P*enf-2* (E). Scale bars are 2 μm. (B, D, F) The diameters of the HIS-72::mCherry^+^ patches shown in (A, C, E) are measured over time and compared to that at “0 min”. The Dt/D0 ratio (DNA degradation index) is plotted over time. The control curves (*nuc-1(e1392)* carrying no transgene) shown in (B), (D), and (F) are from Figure 1 (D), (F), and (H), correspondingly. (G) Histograms of the DNA degradation index of HIS-72::mCherry inside C3 at 48-min post engulfment of *nuc-1(e1392)* mutant with and without the stated transgenes. The numbers of C3s scored for the *nuc-1(e1392)* along, carrying P*ced-1 nuc-1*, and carrying P*efn-2 nuc-1* are 13, 17, and 12, respectively. (H) Fluorescent images showing that a P*efn-2* his-72::mCherry reporter is expressed in C3 but not in its engulfing cells. The co-expressed P*ced-1* PH::GFP reporter labels the plasma membrane of C3’s engulfing cell (ABplaapppp) (yellow arrows). The nucleus of ABplaapppp is labeled with a white arrowhead. C3 is labeled with white arrows. The positions of the two z slices, which are 2 μm apart, are labeled. Scale bars are 2 μm. **Figure 4 ALT TEXT:** Images and graphs related to *nuc-1* rescue in engulfing and dying cells, with subfigures labelled from a to h showing NUC-1 activity by tracking the clearance or lack of clearance of an mCherry-tagged histone reporter.

Together, these results indicate that NUC-1 acts in the engulfing, but not dying, cell for the degradation of dying cell chromatin DNA.

### NUC-1 is not observed inside the nuclei of apoptotic cells

If, as proposed, NUC-1 acts in the nuclei of apoptotic cells to degrade chromatin DNA, one would expect to observe NUC-1 inside the apoptotic cell nuclei. We expressed a NUC-1::mCherry reporter under the control of the *his-72* promoter (P*his-72*), a promoter that is expressed in all cells in embryos, including cells that undergo apoptosis (Ooi, Priess, and Henikoff 2006). We co-expressed this reporter with an integrated nuclear envelope reporter GFP::NPP-22 (Huelgas-Morales et al. 2020). In all the cells observed, NUC-1::mCherry was localized to puncta in the cytoplasm, consistent with its documented lysosomal localization (Fig. 5 A-C) (Guo et al. 2010). On the contrary, no obvious nuclear localization of the mCherry signal was observed in the apoptotic cell Cm before it was engulfed (Fig. 5 A, D, E) or in live cells (Fig. 5 B, D, E). Similarly, in the *ced-5(n1812)* mutant background, no mCherry signal was observed in the Cms that remained unengulfed (Fig. 5 C, D, E). On the other hand, the mCherry signal was observed to accumulate inside the phagosomes after Cm is engulfed (Fig. 5 A, D, E). These results indicate that there is no detectable NUC-1 inside the nuclei of apoptotic or live cells.

**Figure 5.**
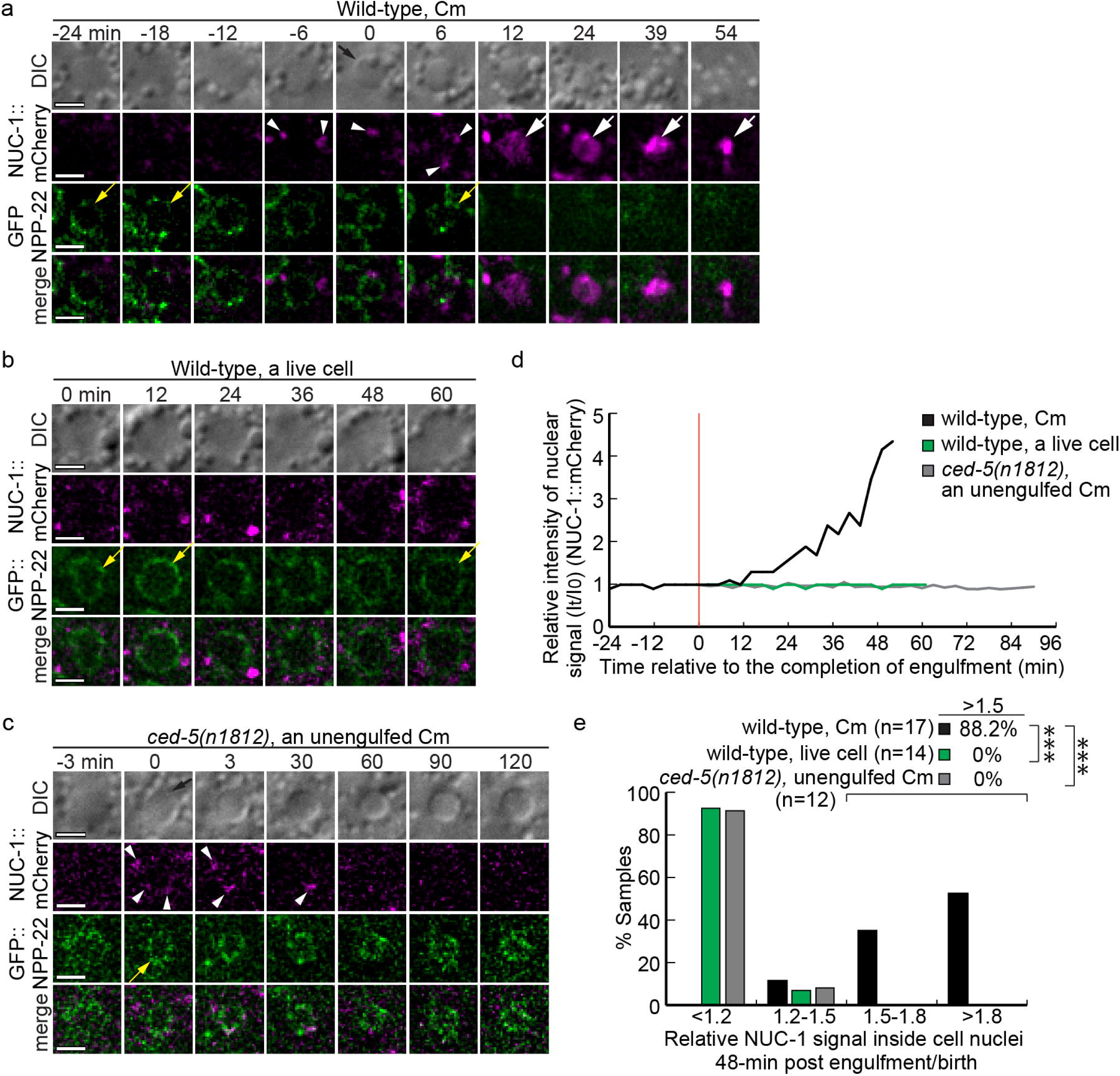
NUC-1 is not localized to the nuclei of apoptotic or live cells. (A-C) Time-lapse images monitoring the subcellular localization of NUC-1::mCherry expressed in all cells as puncta under the control of P*his-72* (white arrowheads) and the corresponding nuclear envelope reporter (GFP::NPP-22, yellow arrows) in Cm (A, C) and a live cell in the same embryos (B). “0 min” in (A, C) is the time point when the button-like DIC morphology first appeared, in (B) is when the live cell was born. In (A), “0 min” is considered the completion of engulfment. The big white arrows in (A) indicate phagosomal lumen filled with NUC-1::mCherry. Scale bars are 2 μm. (D) Curves showing the relative NUC-1::mCherry signal intensity (Intensity(t)/Intesity(t0)) at the center of the nucleus over time. The red vertical line indicates the “0 min” time point. The ratio 1.0 means no significant NUC-1 in the nucleus. The source of samples is from A-C. (E) Histograms of the relative NUC-1::mCherry signal intensity in the nuclei of engulfed or unengulfed Cm and in live cells in two different genetic backgrounds 48 min-post engulfment or birth of the live cells. “***”: p value < 0.001, Student *t*-test, comparing with Cms in wild-type embryos. **Figure 5 ALT TEXT:** Images and graphs related to NUC-1 localization, with subfigures labelled from a to e tracking NUC-1 enrichment along with a nuclear membrane marker.

The optimal pH value for NUC-1 activity is reported to be between 4-5 (Hevelone and Hartman 1988). Previously, using a P*his-72 his-72::gfp::mCherry* reporter and taking advantage of the acid-sensitive and acid-resistant features of GFP and mCherry, respectively, we established an assay that has demonstrated that the phagosomal lumen is gradually acidified until the GFP signal disappears (Peña-Ramos et al. 2022; Peña-Ramos and Zhou 2023) (Fig. S4 C and E).

Using the same reporters, we measured the acidification status of cells destined for apoptosis. We found that before being engulfed, the acidification status (AS) of Cm (Fig. S4A, yellow arrowheads, B) (**Materials and Methods**), which is represented by the intensity ratio between the GFP and mCherry signals, is similar to that of a live neighboring cell (Fig. S4A, yellow arrowheads, B). The nuclei of live cells are known to hold a neutral pH (Casey, Grinstein, and Orlowski 2010). Our results thus indicate that, like live-cell nuclei, cells programmed to undergo apoptosis maintain neutral nuclear pH values before engulfment, which are not favorable to the activity of DNase II endonucleases. Together, the results shown in Figs. 5 and S4 indicate that NUC-1 is unlikely to act inside the apoptotic cell nuclei to initiate chromatin DNA degradation.

### Lysosome-phagosome fusion is critical for the degradation of apoptotic cell DNA

NUC-1 was reported to reside in lysosomes (Guo et al. 2010). By co-expressing NUC-1::mCherry and CTNS-1::GFP (a lysosomal membrane protein (Yu et al 2006)), both under the control of P*his-72* (Fig. S5C) or P*ced-1* (Fig. S5F), we confirmed that they are co-localized to lysosomal puncta. In time-lapse recording experiments, they are both enriched on phagosomal surfaces after the formation of a nascent phagosome (Fig. S5F). Over time, NUC-1::mCherry enters the phagosomal lumen, whereas CTNS-1::GFP remains on the phagosomal surfaces (Fig. S5F). The lysosomal localization of NUC-1 is dependent on its N-terminal Signal Sequence (aa 1-21) (Fig. S5A). Shielding this motif with an N-terminal mCherry fusion (mCherry::NUC-1) or deleting the Signal Sequence (mCherry::ΔssNUC-1) both eliminate the puncta morphology of NUC-1 and its entry into the phagosomal lumen (Fig S5. D, E, G, H). In addition, NUC-1::mCherry expressed under the control of the *nuc-1* promoter (P*nuc-1*) is broadly present in many cells as puncta (Fig. S5B), indicating that NUC-1 resides in the lysosomes of most, if not all, cells.

RAB-7 is a small GTPase that promotes lysosome-phagosome and autophagosome-phagosome fusions in *C. elegans*, as well as fusions between other organelles (Yu, Lu, and Zhou 2008; Peña-Ramos et al. 2022). We observed that in the *rab-7(ok511)* null mutant embryos, the NUC-1::mCherry puncta produced in engulfing cells failed to accumulate inside phagosomes (Fig. 6 A-D). Furthermore, in *rab-7(ok511)* mutants, the time-lapse recording assay monitoring the HIS-72:: mCherry^+^ patches revealed a lack of chromatin DNA degradation in engulfed Cm (Fig. 6 E-G). More broadly, this severe apoptotic cell DNA degradation defect was also observed in many cell corpses in 1.5-fold-stage *rab-7(ok511)* mutant embryos using the HIS-72::mCherry reporter, where cell corpses are distinguished by the button-like DIC morphology (Fig. 6 H-I). The specific expression of *rab-7* in engulfing cells under the control of P*ced-1* rescued the DNA degradation defect of *rab-7(ok511)* mutants not only in Cm (Fig. 6 F-G), but also other embryonic apoptotic cells (Fig. 6 H-I), indicating efficient rescue. Together, these results indicate that the fusion of lysosomes to phagosomes driven by RAB-7, which occurs in engulfing cells, is essential for the delivery of NUC-1 into the phagosomal lumen.

**Figure 6.**
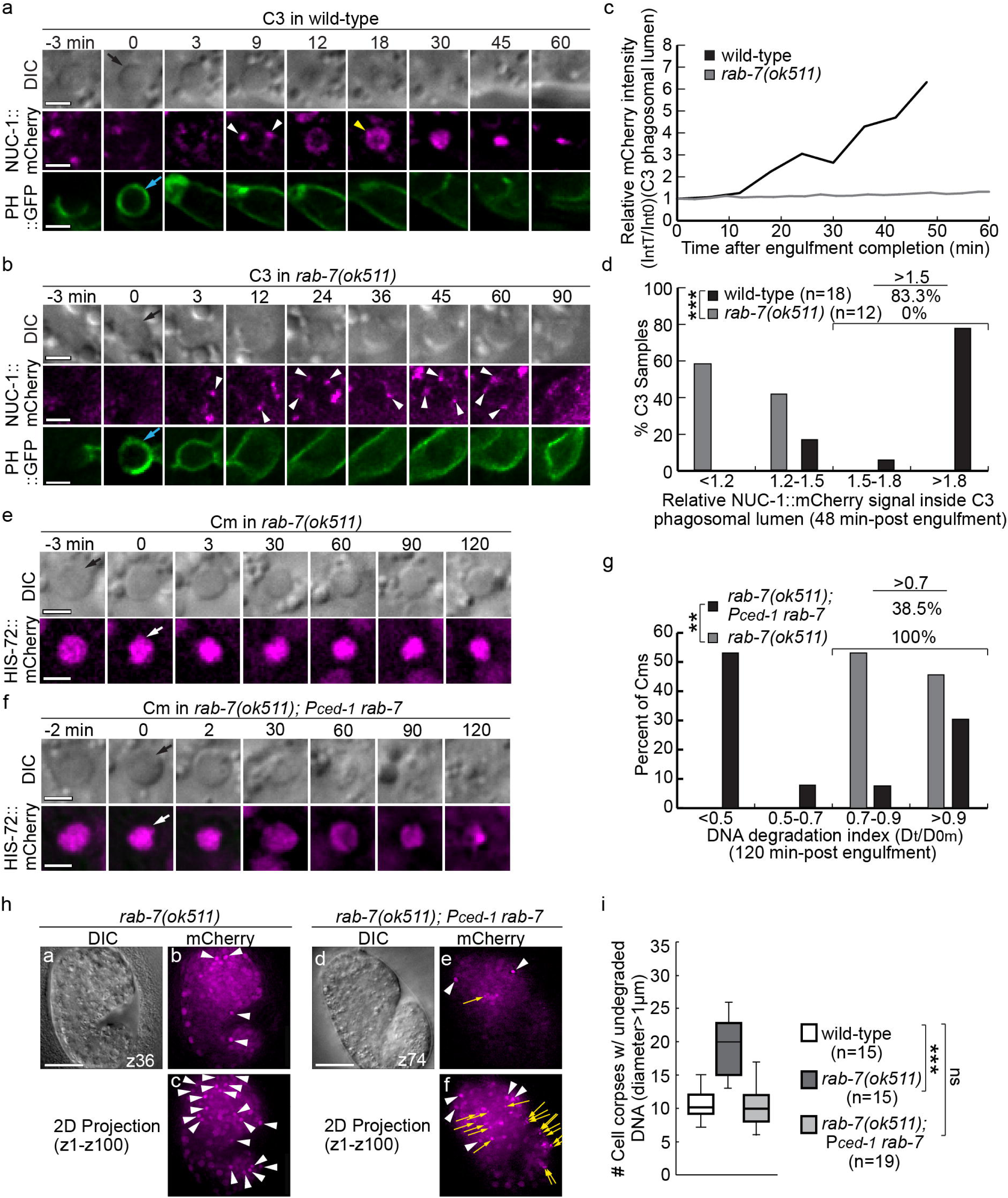
The function of RAB-7 in engulfing cells is essential for the delivery of NUC-1::mCherry into the phagosomal lumen and the DNA degradation of engulfed apoptotic cells (A-B) Time-lapse recording images monitoring the recruitment of NUC-1::mCherry puncta (white arrowheads) produced under P*ced-1* to phagosomal surfaces and its subsequent entry into the phagosomal lumen in C3-containing phagosomes in wild-type (A) and *rab-7(ok511)* mutant (B) embryos. The co-expressed P*ced-1* PH::GFP reporter marks extending pseudopods and enables the determination of the moment when the nascent phagosomes (blue arrows) just form, the “0 min” moment. Black arrows mark the button-like appearance of the C3 cell corpse in the DIC channel. Yellow arrowheads label phagosomes filled with NUC-1::mCherry. Scale bars are 2 μm. (C) The relative intensity (IntT/Int0) of the NUC-1::mCherry signal in the center of the C3 phagosome in wild-type (A) and *rab-7(ok511)* mutant (B) images over time. (D) Histograms of the relative NUC-1::mCherry signal intensities (IntT/Int0) in the center of the C3 phagosomal lumen in wild-type and *rab-7(ok511)* mutant embryos 48 min-post engulfment. “***”, p< 0.001, Student *t*-test. n, number of C3s analyzed. (E-F) Time-lapse recording images monitoring the shrinkage of the HIS-72::mCherry patches (white arrows) inside Cm in *rab-7(ok511)* mutant (E) and *rab-7(ok511)* embryos carrying P*ced-1 rab-7* (F). “0 min”, the moment engulfment is just completed. Black arrows mark Cm when the button-like DIC morphology first arose. Scale bars are 2 μm.| (G) Histograms of the DNA degradation index of the chromatin DNA inside engulfed Cm in *rab-7(ok511)* mutants carrying or not carrying P*ced-1 rab-7*. The data are from time-lapse recording series of 13 Cms in each strain. “**”, 0.001 < p < 0.01, Student *t*-test. (H) DIC and mCherry images of 1.5-fold stage *rab-7(ok511)* embryos carrying P*ced-1 rab-*7 or not that also carried P*his-72 his-72::mCherry* are shown. Cell corpses (distinguished by the DIC morphology) with undegraded chromatin DNA (HIS-72::mCherry diameters >1.0 μm) (white arrowheads) were counted for (I). (a-b, and d-e) are individual z-slices. (c and f) are 2D projection images of the entire Z-stacks of the embryos (100 z-slices at 0.2-μm z-thickness). Yellow arrows mark the remnant of chromatin DNA of engulfed apoptotic cells after degradation. Scale bars are 15 μm. (I) Box-whiskers plots summarizing the quantification of the DNA degradation efficiencies of 1.5-fold stage wild-type embryos and *rab-7(ok511)* embryos carrying P*ced-1 rab-7* or not. “***”: p<0.001, “ns”: not significant (p>0.05), Student *t*-test. n, the number of embryos analyzed. **Figure 6 ALT TEXT:** Images and graphs related to the effect RAB-7 has on cell corpse DNA clearance, with subfigures labelled from a to i illustrating lysosome recruitment and fusion as well as cell corpse clearance in wild-type and rab-7 null mutant backgrounds.

In addition, we found that *rab-7* is essential for phagosome acidification in embryos, most likely through regulating lysosome-phagosome fusion. In *rab-7(ok511)* mutant embryos, time-lapse imaging of phagosomes containing Cm revealed that acidification is severely deficient (Fig. S4 C-F), indicating that *rab-7* is required for multiple phagosome maturation events.

### The delivery of NUC-1 into phagosomes is regulated by the CED-1 phagocytic receptor

Previously, we have reported that the phagocytic receptor CED-1 is essential for promoting the recruitment and fusion of lysosomes and other intracellular organelles to phagosomes (Yu, Lu, and Zhou 2008; Peña-Ramos et al. 2022). To further examine whether the entry of lysosomal components into the phagosome is CED-1-dependent, we monitored the dynamic localization pattern of the NUC-1::mCherry puncta expressed in engulfing cells under P*ced-1* (Fig. 7 A-C). We first measured the time when the NUC-1::mCherry puncta enrichment on the surfaces of phagosomes containing the apoptotic cell C5 started (Fig. 7D), an indication of recruitment of lysosomes. We found that the loss of *ced-1* function does not affect the timing when enrichment started (Fig. 7D). However, the mCherry signal on the phagosomal surfaces and inside the phagosomal lumen is much reduced in *ced-1(e1735)* null mutants (Fig. 7 B, C, and F). Quantitation of the mCherry relative signal intensity at the center of the phagosomal lumen further demonstrates that the entry of NUC-1::mCherry into the lumen is significantly delayed in *ced-1* mutants (Fig. 7E); furthermore, mCherry accumulation inside the phagosomal lumen is greatly reduced (Fig. 7F). These results indicate that CED-1 indeed is the upstream regulator that drives the delivery of NUC-1 and possibly other lysosomal enzymes into the phagosomal lumen.

**Figure 7.**
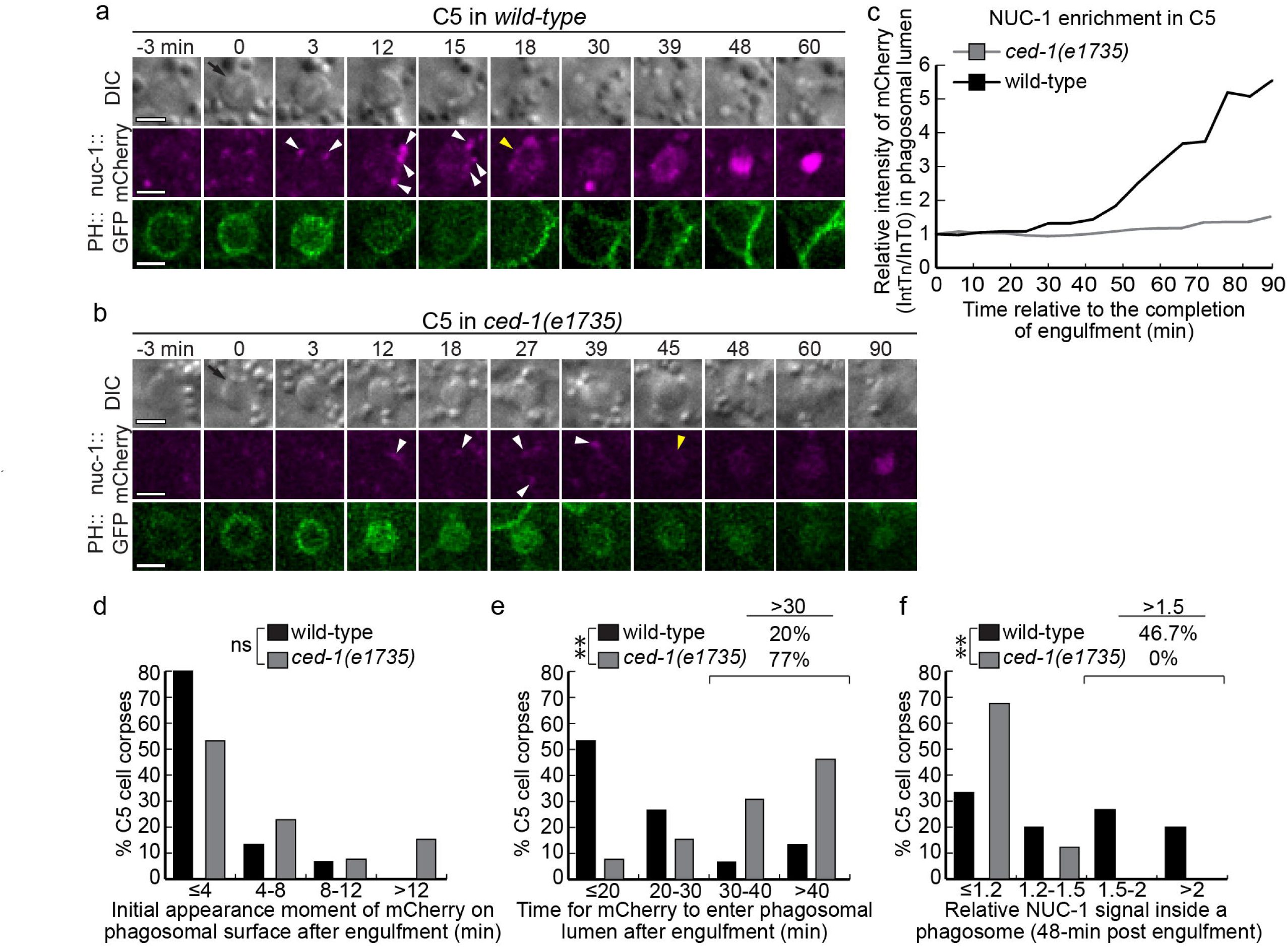
In *ced-1* mutants, the delivery of NUC-1 into the phagosomal lumen is severely impaired. (A-B) Time-lapse images monitoring NUC-1::mCherry expressed in engulfing cells under P*ced-1*, the co-expressed pseudopod marker P*ced-1* PH::GFP, and the DIC morphology of C5 (black arrows) before and after the completion of engulfment (“0 min”), indicated by the newly sealed GFP ring. White arrowheads label NUC-1::mCherry puncta recruited to the phagosomal surface. Yellow arrowheads indicate phagosomal lumen filled with NUC-1::mCherry. Scale bars are 2 μm. (C) Curves showing the relative NUC-1::mCherry signal intensity (Int(Tn)/Int(T0)) at the center of the phagosome over time. The “0 min” time point is the completion of engulfment. The ratio 1 means no significant NUC-1 in the phagosome.The data are from (A) and (B). (D-F) Histograms illustrating the recruitment of the NUC-1::mCherry puncta to phagosomal surfaces (D), the time that NUC-1::mCherry is 1^st^ detected in the phagosomal lumen (E), and the relative mCherry intensities (Int(T48)/Int(T0) in the center of the phagosomal lumen (F) of 15 C5 in wild-type embryos and 13 *ced-1(e1735)* mutant embryos. **Figure 7 ALT TEXT:** Images and graphs related to the effect CED-1 has on cell corpse DNA clearance, with subfigures labelled from a to f illustrating NUC-1 localization and enrichment in wild-type and *ced-1* null mutant backgrounds.

## Discussion

*nuc-1* was the first *C. elegans* gene identified that acts in the clearance of apoptotic cells. This report aims to determine which of the two contradictory models about the molecular mechanism of NUC-1 action in the literature is true. Model 1 proposes that NUC-1 acts in the lysosomes of engulfing cells to degrade the chromatin DNA of engulfed apoptotic cells. Model 2, on the other hand, proposes that NUC-1 acts in the apoptotic cells, inside the nuclei, to digest the chromatin DNA cell-autonomously. Our findings indicate that Model 2 is unlikely to be true. First of all, based on the observation that in engulfment mutants (*ced-2*, *ced-5*, *ced-6*, and *ced-10*), chromatin DNA was still degraded, and based on the assumption that in engulfment mutants, all apoptotic cells remained unengulfed, Wu, Stanfield, and Horvitz (2000) concluded that the function of NUC-1 in degrading chromatin DNA did not require engulfment. The same approach also plays an essential role in reaching the same conclusion by Lai et al. (2009), Yu et al. (2015), and Aruscavage, Hellwig, and Bass (2010). For example, by comparing the number of ToLFP-positive spots between wild-type and *ced-1* mutants, which is presumed to be engulfment-deficient, Yu et al. (2015) conclude that the DNase II activity is 70% autonomous in apoptotic cells and 30% non-autonomous from other cells. On the contrary, through monitoring the fate of apoptotic cells via time-lapse recording in the *ced-1*, *ced-2*, and *ced-5* loss-of-function mutants, we found that in all of these mutants, engulfment was only partially defective, and substantial percentages of apoptotic cells were engulfed (Fig. S3A). For example, 94.7%, 80.0%, and 79.1% of C3 are engulfed in *ced-1*, *ced-2*, and *ced-5* loss-of-function mutants, respectively (Fig. S3A). Therefore, the 1.5-fold stage engulfment mutant embryos analyzed by Wu, Stanfield, and Horvitz (2000), Lai et al. (2009), Yu et al. (2015), and Aruscavage, Hellwig, and Bass (2010) actually contained a mixture of engulfed and unengulfed apoptotic cells. As a result, many cell corpses that had been presumed to be unengulfed were actually engulfed.

To avoid the above mis-presumption, we thus analyzed chromatin DNA degradation in unengulfed and engulfed apoptotic cells separately via time-lapse recording and a “DNA degradation assay in whole embryos” that was able to distinguish engulfed cell corpses from other cells. Our results clearly indicate that the chromatin DNA of unengulfed apoptotic cells remains undegraded, yet that of engulfed apoptotic cells is properly degraded despite the *ced-5(ok511)* null mutation (Fig. 3). These results indicate that being engulfed is required for the degradation of apoptotic chromatin DNA to occur. In support of this conclusion, we further demonstrated that the specific expression of *nuc-1* in engulfing, but not dying, cells efficiently rescued the DNA-degradation defect of the *nuc-1(e1392)* mutants. Together, our results demonstrated that *nuc-1* acts in engulfing cells to degrade the chromatin DNA of engulfed apoptotic cells.

Another piece of evidence that casts doubt on any role of NUC-1 as an endonuclease acting inside the apoptotic cell nuclei comes from the biochemical property of NUC-1. Like DNase II, its mammalian homolog, NUC-1 is only active under acidic conditions (Hevelone and Hartman 1988; Evans and Aguilera 2003). NUC-1 is inactive in an *in vitro* DNA degradation assay when the pH is higher than 6.2, and the optimal pH for its activity is 4-5 (Hevelone and Hartman 1988). The nuclei of eukaryotic cells hold a pH value around 7.2 (Casey, Grinstein, and Orlowski 2010), at which NUC-1 should be inactive. In *C. elegans* embryos, we further measured the acidification status of the nuclei of apoptotic cells, a representation of their pH values, at time points before being engulfed, and found that the acidification status of these nuclei is similar to that of the nuclei of the neighboring living cells, which is unfavorable for NUC-1 activity.

Previously, whether NUC-1 was located inside the nucleus was not reported. A striking piece of evidence we have observed is that in the nuclei of living cells or apoptotic cells before they are engulfed, no NUC-1::mCherry is detected. Previously, Lai et al. (2009) suggested that NUC-1 is a secreted protein that is produced in other cell types and is subsequently endocytosed into apoptotic cells and enters the nuclei to degrade chromatin DNA. However, Lai et al. (2009) did not report any evidence of the nuclear localization of NUC-1. This hypothesis does not agree with the common understanding of how endocytosed cargos travel along the endolysosomal system (Cullen 2008). Our failure to detect NUC-1::mCherry in the nuclei of living or apoptotic cells further disagrees with this hypothesis. Together, the requirement of engulfment for DNA degradation, the finding that NUC-1 acts in engulfing but not in dying cells, the finding that, before being engulfed, apoptotic cell nuclei maintain a neutral pH, and the lack of detection of NUC-1 inside nuclei strongly indicate that NUC-1 does not act cell autonomously to degrade apoptotic chromatin DNA inside the nucleus.

In addition to demonstrating that Model 2 is not likely to occur, we also further provided lines of evidence that support Model 1 and bring it more molecular details. Using time-lapse imaging and a fluorescent pH reporter, we report the gradual acidification of the phagosomal lumen. Previously, Guo et al. (2010) reported that NUC-1 is localized to the lysosomes and reported three states of NUC-1::mCherry in relation to the phagosome: “attaching”, “clustering”, and “incorporated”. We further observed the sequential events of the recruitment and further enrichment of the NUC-1::mCherry puncta to the phagosomal surfaces, followed by the entry of the NUC-1::mCherry signal into the phagosomal lumen, corresponding to the “attaching”, “clustering”, and “incorporated” status, respectively. These are the events occurring during lysosome-phagosome fusion. Furthermore, we discovered that the delivery of NUC-1 into the phagosomal lumen and the degradation of apoptotic chromatin DNA both required lysosome-phagosome fusion inside engulfing cells, again demonstrating that lysosomes in engulfing cells are the source of the NUC-1 endonuclease. Previously, we reported that the phagosomal lumen is acidified (Peña-Ramos et al. 2022). In this report, we further find that the acidic phagosomal pH requires lysosome-phagosome fusion (Fig. S4). This acidic environment is suitable for the enzymatic activity of NUC-1. Our findings thus strongly support the conclusion that NUC-1 acts in the lysosomal lumen. Last but not least, in this report, we observe that the phagocytic receptor CED-1 drives the delivery of NUC-1 into the phagosomal lumen. This result is consistent with our previous finding that CED-1 controls the recruitment of lysosomes and other intracellular organelles to the phagosomal surfaces and the subsequent fusion of these organelles to phagosomes (Yu, Lu, and Zhou 2008; Peña-Ramos et al. 2022). NUC-1 is the first *C. elegans* lysosomal hydrolase discovered that specifically targets components of engulfed apoptotic cells.

We found that the predicted Signal Sequence (ss) is necessary for the localization of NUC-1 to lysosomes. Deleting or shielding this Signal Sequence results in the cytoplasmic localization of NUC-1. Besides being a lysosomal luminal protein, NUC-1 is also required for the degradation of bacterial DNA inside the intestinal lumen (Hevelone and Hartman 1988; Lai et al. 2009; Yu et al. 2015) and is proposed to be a secreted protein. How it is determined whether NUC-1 is secreted or retained in the lysosome of a given cell is a challenging question for future study.

In mice, lysosomal acidification in macrophages was reported to be required for the degradation of the DNA of engulfed apoptotic cells (McIlroy et al, 2000). Mammalian DNase II is a lysosomal enzyme and an essential gene. During erythropoiesis, erythroblasts undergo enucleation to become erythrocytes. The expelled nuclei are engulfed by macrophages, and the chromatin DNA is digested by DNase II in the macrophages. In DNase II -/-mouse embryos, the erythroblast chromatin DNA inside phagosomes is not degraded, resulting in the blockage of erythroblast enucleation and causing severe anemia and embryonic lethality (Kawane et al. 2001). In addition, the DNA of embryonic apoptotic cells in the thymus remains undegraded, activating innate immunity and leading to defects in thymic development (Kawane et al. 2003). Conditional deletion of DNase II in adult mice blocks the degradation of mammalian DNA from erythroid precursors and apoptotic cells, leading to the production of proinflammatory cytokines and the development of chronic polyarthritis resembling human chronic arthritis (Kawane et al. 2006). These results indicate that degrading phagosomal cargos in a cell non-autonomous manner has a profound impact on anti-inflammation. Our findings reported here strongly indicate that NUC-1, like mammalian DNase II, also acts cell non-autonomously to degrade the DNA of engulfed apoptotic cells. Therefore, an evolutionarily conserved molecular mechanism governs the DNA degradation event.

## Data availability statement

The authors affirm that all data necessary for confirming the conclusions of the article are present within the article, figures, and tables. Strains and plasmids are available upon request.

## Acknowledgments

We thank Nicole Auld for technical support. We thank Dr. Kota Mizumoto for the pSM-mStayGold plasmid and the *C. elegans* Genetics Center (CGC), funded by the NIH Office of Research Infrastructure Programs (P40 OD010440), for providing some strains. We also thank Dr. H. R. Horvitz for the *nuc-1(e1392)* mutant strain.

## Study Funding

| Funder | Grant Reference Number | Author |
| --- | --- | --- |
| NIH | R01GM067848 | Zheng Zhou |
| NIH | R35 GM153415 | Zheng Zhou |
| NIH | R01NS129847 | Zheng Zhou |

The funders had no role in study design, data collection and interpretation, or the decision to submit the work for publication.

## Supplemental Figures

**Figure S1.**
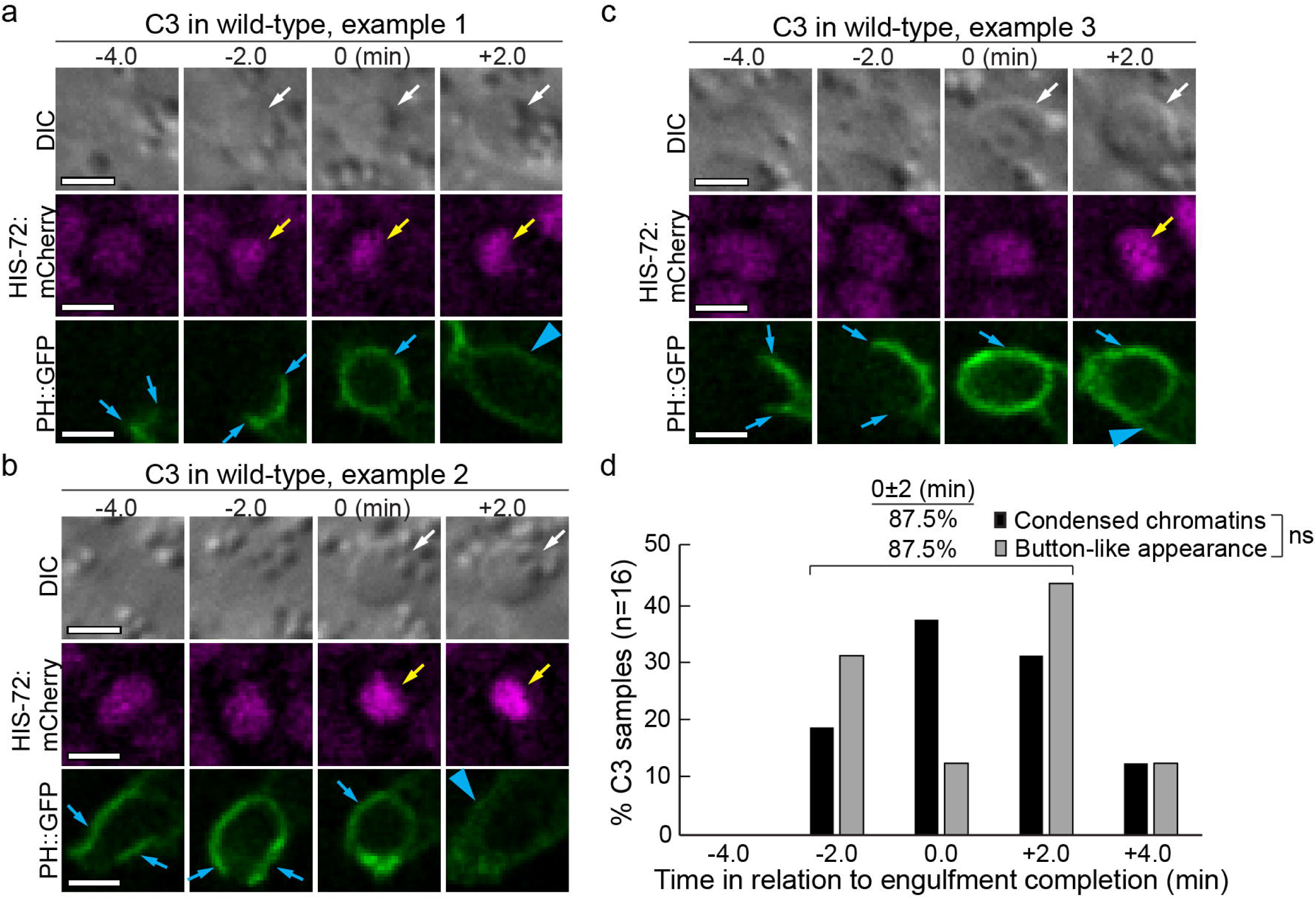
In wild-type embryos, chromatin condensation and the rise of the button-like morphology of an apoptotic cell coincide with the completion of its engulfment. (A-C) The time-lapse series of three C3 apoptotic cells around the time of the completion of their engulfment (0 min). White arrows indicate the C3 apoptotic cells that have assumed a button-like morphology under the DIC optics. The pseudopods from the engulfing cell (blue arrows) and the nascent phagosomes containing C3 (blue arrowheads) are labeled with the P*ced-1* PH(PLCγ)::GFP (PH::GFP) expressed only in engulfing cells. The condensed chromatin of C3 (yellow arrows) is visualized by the HIS-72::mCherry reporter. Scale bars are 2 μm. (D) A histogram displaying the distribution of the time points when the button-like appearance of C3 starts to rise and when the condensation of the chromatin of C3 starts to be detected. The number of C3s analyzed is 16. **Figure S1 ALT TEXT:** Images and graph related to phenotypic markers that are temporally correlated with phagocytic engulfment of apoptotic cell corpses, with subfigures labelled from a to d illustrating 3 different temporal sequences of these phenotypes.

**Figure S2.**
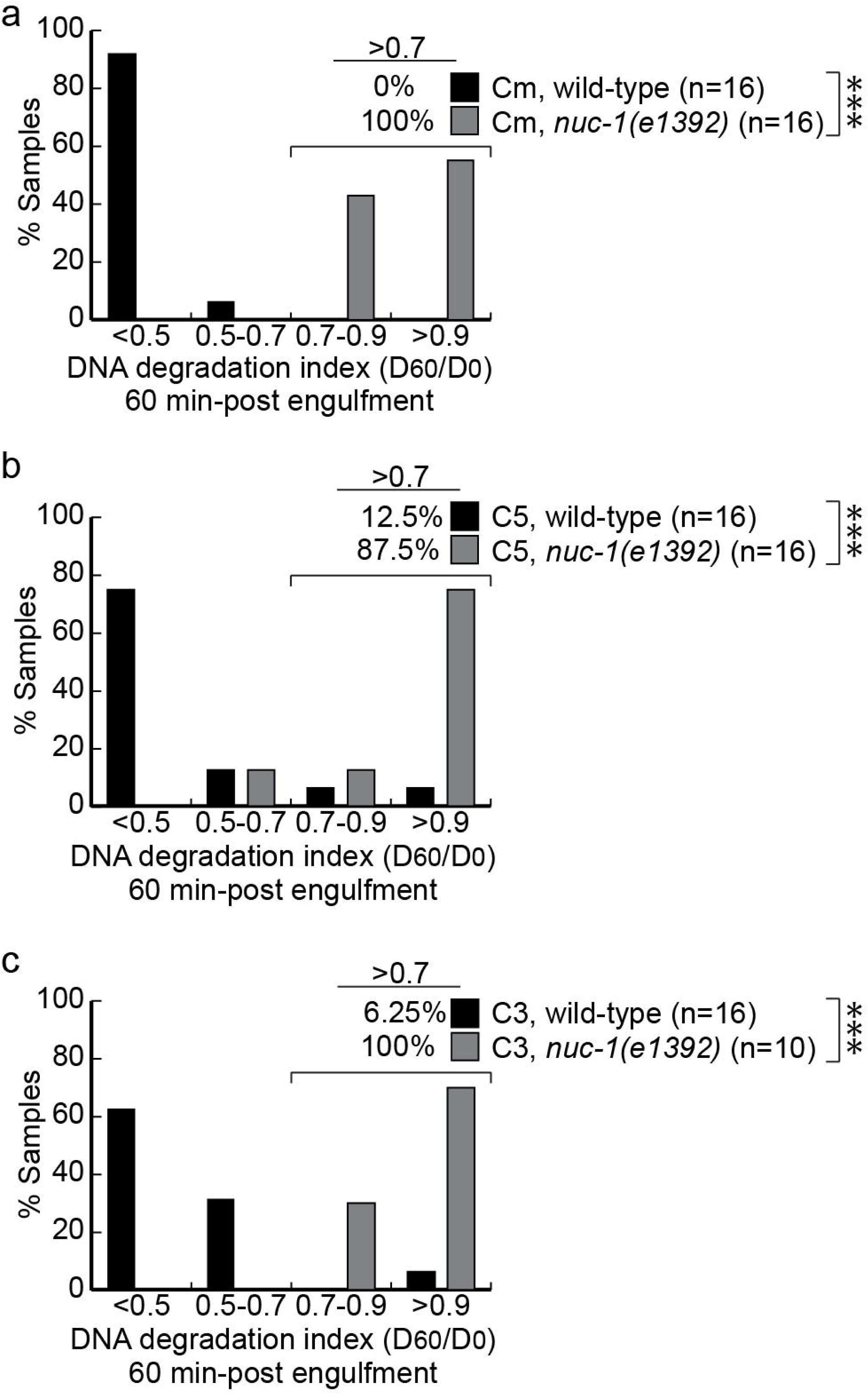
Histograms of quantitative DNA degradation index of multiple Cm, C5, and C3s as shown in Figure 1 at 60 min-post engulfment completion. (A), (B), and (C) correspond to the same analysis (C), (E), and (G), of Figure 1, were obtained from. n, the number of samples analyzed. “***”, p<0.001. p-value is from Student *t*-test. **Figure S2 ALT TEXT:** Graphs showing the proportional distribution of data from Figure 1, with subfigures labelled from a to c corresponding to subfigures in Figure 1.

**Figure S3.**
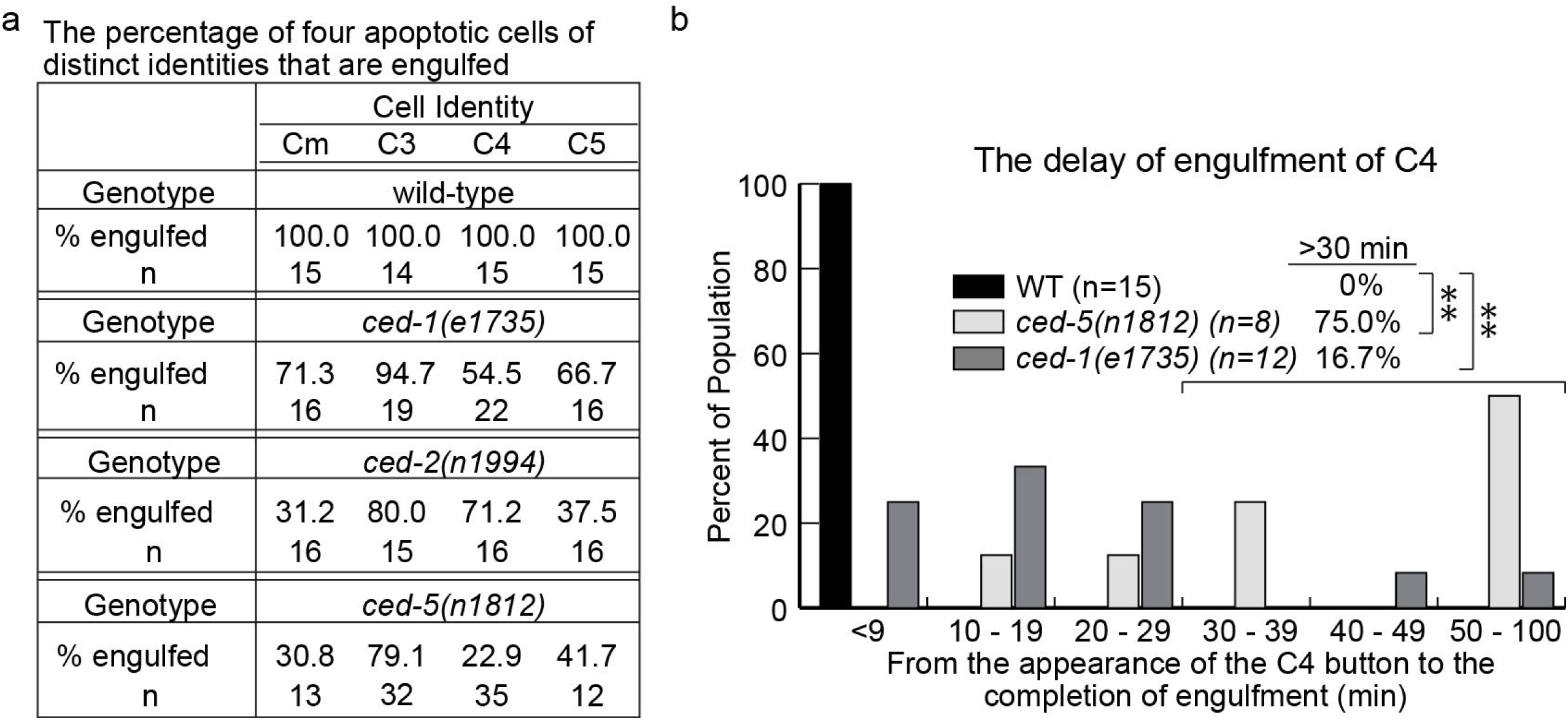
Time-lapse recording images reveal that in *ced-1*, *ced-2*, and *ced-5* mutants, certain percentages of apoptotic cells are engulfed. (A) A table quantifying whether apoptotic cells Cm, C3, C4, and C5 are engulfed in multiple embryos of different genotypes. The time point “0 min” is defined as the moment when the button-like DIC morphology first appears, which indicates apoptosis. If, after 100 min, the PH(PLCγ)::GFP-labeled pseudopods from engulfing cells still haven’t successfully enclosed a dying cell, the dying cell is considered unengulfed. n, the number of embryos scored. (B) A histogram, among C4s that have been engulfed, of the time it takes to engulf C4 in embryos of different genotypes, determined by the extension and seal of the PH(PLCγ)::GFP-labeled pseudopods. n, the number of embryos scored. “**”, 0.001 < p < 0.01, Student *t*-test. **Figure S3 ALT TEXT:** Table and graph related to the penetrance of engulfment mutants, with subfigures labelled from a and b detailing engulfment totals and delays in engulfment.

**Figure S4.**
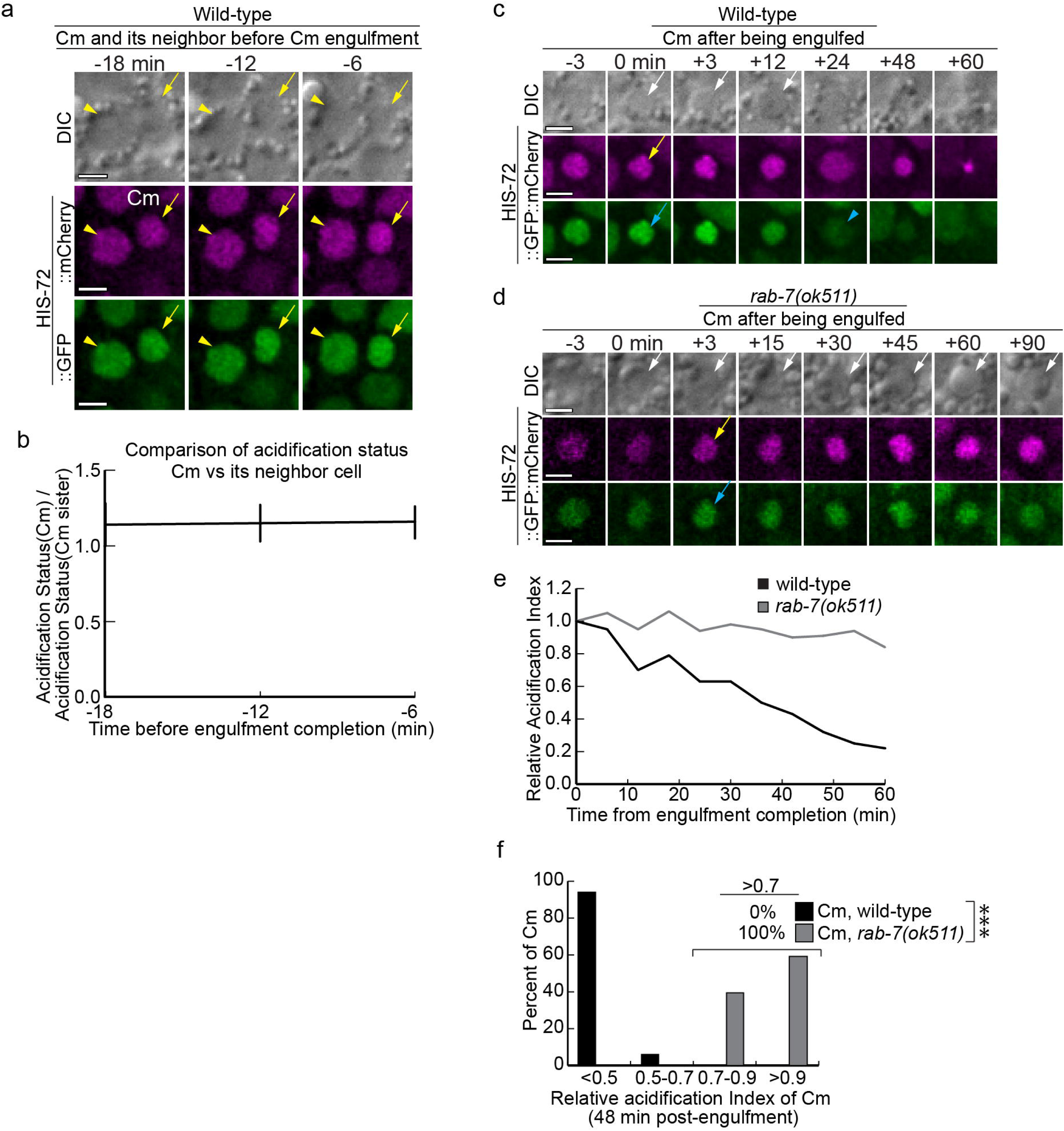
The nucleus of an apoptotic cell is not acidified until it is engulfed inside a phagosome. In addition, *rab-7* mutants are defective in the acidification of the phagosomal lumen. (A) The images of the GFP and mCherry signals in Cm (yellow arrows) and a nearby cell (yellow arrowheads) in a wild-type embryo carrying P*his-72 his-72::gfp::mCherry*. Time points label the moments before the completion of engulfment (the “0 min” time point). Scale bars are 2 μm. (B) Comparison of the acidification status between Cm and the nearby cell at the same time points as in (A). The numbers on the Y-axis are (IGFP/ImCherry)Cm / (IGFP/ImCherry)Cm sister. Error bars indicate standard deviation (sd). 13 embryos were scored.| (C-D) Time-lapse images monitoring the GFP and mCherry signal at the center of an engulfed Cm (white arrow) in wild-type (C) and *rab-7(ok511)* mutant (D) embryos expressing P*his-72 his-72::gfp::mCherry*. The “0 min” time point is when the button-like DIC morphology (white arrows) first arises, which occurs simultaneously with the completion of engulfment. Yellow arrows in the mCherry channel and blue arrows in the GFP channel label nuclei. A blue arrowhead in (C) marks a Cm nucleus in which the GFP signal is quenched. Scale bars are 2 μm. (E) Curves displaying the acidification indices at the center of the Cm nuclei over time in the images displayed in (C) and (D). The relative acidification index (RAI) = (IGFP/ImCherry)Tn / (IGFP/ImCherry)T0. (F) Histogram of the relative acidification index (RAI) of 17 and 10 Cms in the wild-type and *rab-7(ok511)* mutant embryos, respectively, at 48 min post-engulfment. sd: standard deviation. “***”, p<0.001. p-value is from Student *t*-test between the RAI of wild-type and *rab-7(ok511)* samples. **Figure S4 ALT TEXT:** Images and graphs related to acidification of apoptotic cell corpse nuclei, with subfigures labelled from a to f illustrating acidification in apoptotic cell corpses before and after engulfment in wild-type as well as in rab-7 null mutants.

**Figure S5.**
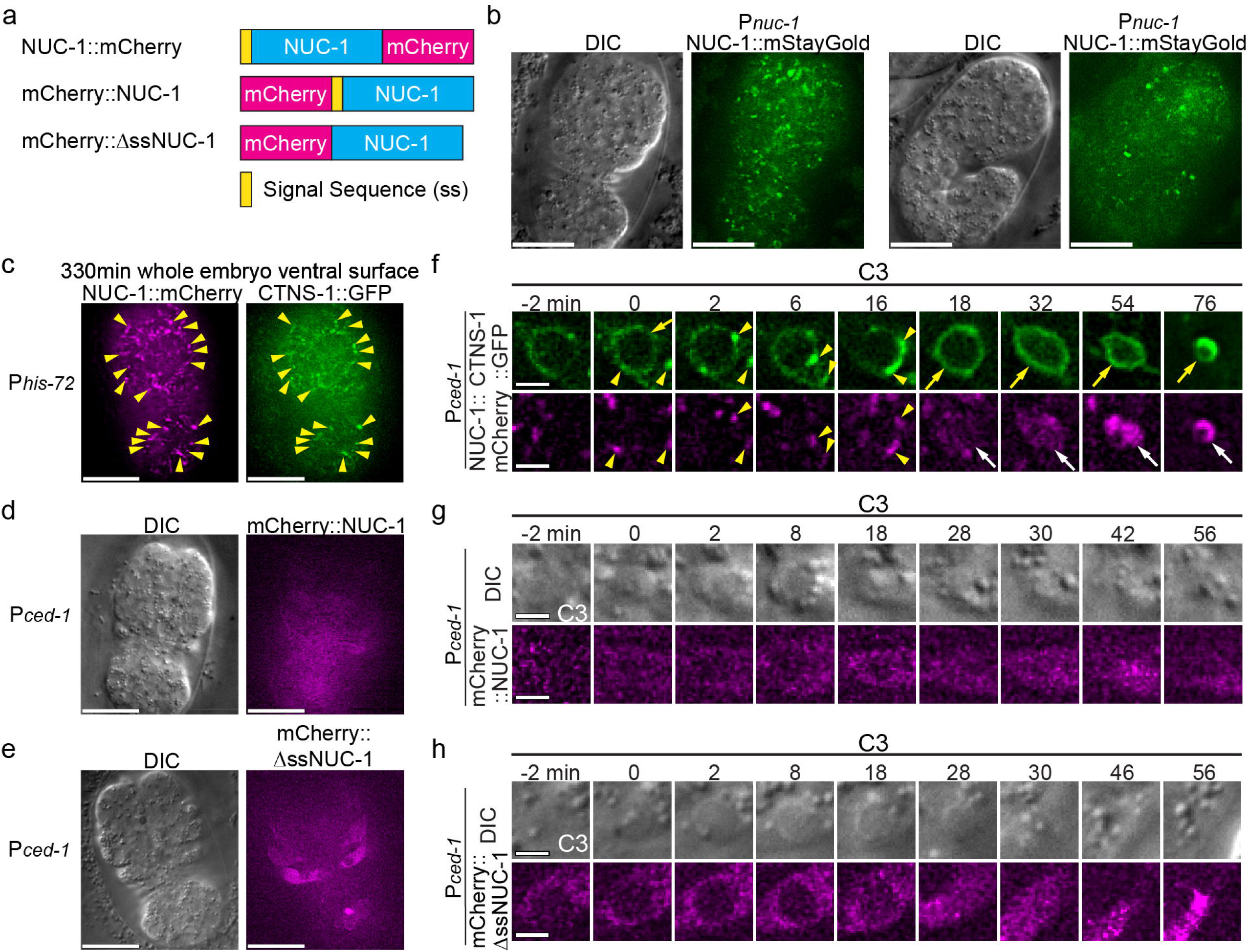
NUC-1 is broadly localized in wild-type embryos and is localized to lysosomes in a manner dependent on its signal sequence (A) Diagrams of the domain structure of NUC-1 and truncated NUC-1 fused to mCherry. (B) The broad localization pattern of NUC-1::mStayGold expressed under P*nuc-1* in two embryos at different stages. (C) In embryos, NUC-1::mCherry colocalizes with CTNS-1::GFP, a lysosomal marker, to intracellular puncta (yellow arrowheads) that are lysosomes. Both reporters are expressed under P*his-72*. (D and E) In embryos, mCherry::NUC-1 and mCherry::ΔssNUC-1 are evenly distributed in the cytoplasm. (B-E) Scale bars are 15 μm. (F, G, and H) Time-lapse images focusing on apoptotic cell C3 and the phagosome C3 is in. “0 min” is the moment when engulfment is just completed. Scale bars are 2 μm. (F) Time-lapse images showing that CTNS-1::GFP and NUC-1::mCherry co-expressed in the engulfing cell for C3 are enriched on the surfaces of phagosomes containing C3, first as puncta (yellow arrowheads); subsequently, the CTNS-1::GFP signal is incorporated into the phagosomal membrane (yellow arrows), yet the NUC-1::mCherry is accumulated in the phagosomal lumen (white arrows). (G-H) mCherry::NUC-1 (G) and mCherry::ΔssNUC-1 (H) expressed in the engulfing cell for C3 are localized in the cytoplasm and not enriched on the phagosomal surfaces throughout the degradation process of C3. **Figure S5 ALT TEXT:** Images related to NUC-1 localization, with subfigures labelled from a to h illustrating cloning strategies to test the effect of promoter and signal sequence changes on NUC-1’s localization pattern in a wild-type background.

