## Supplemental Table 1 for "The *C. elegans* endonuclease NUC-1 acts in engulfing cells to degrade the apoptotic cell DNA"

| Reagent type | Designation | Source or reference | Identifiers | Additional information |
|---|---|---|---|---|
| Strain, strain background *(E. coli)* | OP50 | CGC | OP50 | |
| Strain, strain background *(C. elegans)* | CB911 | CGC | *unc-76(e911) V* | |
| Strain, strain background *(C. elegans)* | CB1392 | CGC | *nuc-1(e1392) X* | |
| Strain, strain background *(C. elegans)* | MT8791 | Horvitz Lab | *ced-5(n1812)IV; unc-76(e911)V* | |
| Strain, strain background *(C. elegans)* | MT9011 | Horvitz Lab | *ced-1(e1735)I; unc-76(e911)V* | |
| Strain, strain background *(C. elegans)* | VC308 | CGC | *rab-7(ok511)II/mIn-1 II* | |
| Strain, strain background *(C. elegans)* | ZH2263 | This study | *enIs71 I; unc-76(e911)V [* $P_{his-72}$ *his-72::mCherry;* $P_{ced-1}$ *PH::gfp;* $P_{unc-76}$ *(+)]* | Figures 1, S1, S2, + S3 Available from the Zhou Lab |
| Strain, strain background *(C. elegans)* | ZH3295 | This study | *enIs71 I; unc-76(e911)V; nuc-1(e1392am)X [* $P_{unc-76}$ *(+);* $P_{his-72}$ *his-72::mCherry;* $P_{ced-1}$ *PH::gfp]* | Figures 1, 2, + S2 Available from the Zhou Lab |
| Strain, strain background *(C. elegans)* | ZH3360 | This study | *unc-76(e911)V; nuc-1(e1392am)X; enEx1723 [* $P_{ced-1}$ *nuc-1::gfp;* $P_{his-72}$ *his-72::mCherry;* $P_{unc-76}$ *(+)]* | Figures 2+4 Available from the Zhou Lab |
| Strain, strain background *(C. elegans)* | ZH3296 | This study | *enIs71 I; ced-5(n1812)IV; unc-76(e911)V [* $P_{unc-76}$ *(+);* $P_{his-72}$ *his-72::mCherry;* $P_{ced-1}$ *PH::gfp]* | Figures 3 + S3 Available from the Zhou Lab |
| Strain, strain background *(C. elegans)* | ZH3823 | This study | *unc-76(e911)V; nuc-1(e1392)X; enEx1961 [* $P_{unc-76}$ *(+);* $P_{efn-2}$ *nuc-1::gfp;* $P_{his-72}$ *his-72::mCherry]* | Figure 4 Available from the Zhou Lab |
| Strain, strain background *(C. elegans)* | ZH3816 | This study | *unc-76(e911)V; nuc-1(e1392)X; enEx1958 [* $P_{unc-76}$ *(+);* $P_{efn-2}$ *his-72::mCherry;* $P_{ced-1}$ *PH::GFP]* | Figure 4 Available from the Zhou Lab |
| Strain, strain background *(C. elegans)* | ZH3735 | This study | *unc-76(e911)V; enEx1866 [* $P_{unc-76}$ *(+);* $P_{his-72}$ *nuc-1::mCherry;* $P_{npp-22}$ *gfp::tev::3xflag::npp-22]* | Figure 5 Available from the Zhou Lab |
| Strain, strain background *(C. elegans)* | ZH3829 | This study | *ced-5(n1812)IV; tn1794; unc-76(e911) V; enEx1869 [* $P_{unc-76}$ *(+);* $P_{npp-22}$ *gfp::tev::3xflag::npp-22;* $P_{his-72}$ *nuc-1::mCherry]* | Figure 5 Available from the Zhou Lab |

| | | | | |
|---|---|---|---|---|
| Strain, strain background *(C. elegans)* | ZH3506 | This study | *enIs71 I; rab-7(ok511)II/mIn-1 II; unc-76(e911)V [$P_{unc-76}$ (+); $P_{his-72}$ his-72::mCherry; $P_{ced-1}$ PH::gfp]* | Figure 6 Available from the Zhou Lab |
| Strain, strain background *(C. elegans)* | ZH3559 | This study | *rab-7(ok511)II/mIn-1 II; unc-76(e911)V; enEx1845 [$P_{unc-76}$ (+); $P_{ced-1}$ GFP::rab-7; $P_{his-72}$ his-72::mCherry]* | Figure 6 Available from the Zhou Lab |
| Strain, strain background *(C. elegans)* | ZH3749 | This study | *rab-7(ok511)II/mIn-1 II; unc-76(e911)V; enEx1158 [$P_{ced-1}$ PH::GFP; $P_{ced-1}$ nuc-1::mCherry; $P_{unc-76}$ (+)]* | Figure 6 Available from the Zhou Lab |
| Strain, strain background *(C. elegans)* | ZH2389 | This study | *unc-76(e911)V; enEx1158 [$P_{ced-1}$ PH::GFP; $P_{ced-1}$ nuc-1::mCherry; $P_{unc-76}$ (+)]* | Figures 6+7 Available from the Zhou Lab |
| Strain, strain background *(C. elegans)* | ZH2386 | This study | *ced-1(e1735)I; unc-76(e911)V; enEx1155 [$P_{ced-1}$ PH::GFP; $P_{ced-1}$ nuc-1::mCherry; $P_{unc-76}$ (+)]* | Figure 7 Available from the Zhou Lab |
| Strain, strain background *(C. elegans)* | ZH2312 | This study | *ced-1(e1735)I; enIs71 I; unc-76(e911)V [$P_{unc-76}$ (+); $P_{his-72}$ his-72::mCherry; $P_{ced-1}$ PH::gfp]* | Figure S3 Available from the Zhou Lab |
| Strain, strain background *(C. elegans)* | ZH760 | This study | *ced-2(n1994)IV; unc-76(e911)V; enIs7 X [$P_{ced-1}$ ced-1::gfp; $P_{unc-76}$ (+)]* | Figure S3 Available from the Zhou Lab |
| Strain, strain background *(C. elegans)* | ZH2059 | This study | *unc-76(e911)V; enEx979 [$P_{his-72}$ his-72::gfp::mCherry; $P_{unc-76}$ (+)]* | Figure S4 Available from the Zhou Lab |
| Strain, strain background *(C. elegans)* | ZH1936 | This study | *unc-76(e911)V; enEx938 [$P_{ced-1}$ ctns-1::GFP; $P_{ced-1}$ nuc-1::mCherry]* | Figure S5 Available from the Zhou Lab |
| Strain, strain background *(C. elegans)* | ZH3627 | This study | *unc-76(e911)V; enEx1886 [$P_{unc-76}$ (+); $P_{his-72}$ nuc-1::mCherry; $P_{his-72}$ ctns-1::GFP]* | Figure S5 Available from the Zhou Lab |
| Strain, strain background *(C. elegans)* | ZH3808 | This study | *unc-76(e911)V; nuc-1(e1392)X; enEx1952 [$P_{unc-76}$ (+); $P_{ced-1}$ mCherry::NUC-1]* | Figure S5 Available from the Zhou Lab |
| Strain, strain background *(C. elegans)* | ZH3817 | This study | *unc-76(e911)V; nuc-1(e1392)X; enEx1959 [$P_{unc-76}$ (+); $P_{ced-1}$ mCherry::(Δss)NUC-1]* | Figure S5 Available from the Zhou Lab |
| Strain, strain background *(C. elegans)* | ZH3847 | This study | *unc-76(e911)V; enEx1972 [$P_{unc-76}$ (+); $P_{nuc-1}$ nuc-1::mStayGold]* | Figure S5 Available from the Zhou Lab |

| | | | | |
|---|---|---|---|---|
| Oligo | ZZ1036 | This study | 5'-aacccGCATGCggacaaggtggacgagaaatcc-3' | Available from the Zhou Lab |
| Oligo | ZZ1037 | This study | 5’-GGTTTGGATCCAGCGGCAGGAGACAAGCCCAT-3' | Available from the Zhou Lab |
| Oligo | ZZ1227 | This study | 5'-aacccgcatgcgtgtggacaccaatttggc-3' | Available from the Zhou Lab |
| Oligo | ZZ1228 | This study | 5'-ggcaaggatcctgttgttctggaaattgagaat-3' | Available from the Zhou Lab |
| Oligo | ZZ1268 | This study | 5’-AAGCTTGCATGCgtgtggacacc-3’ | Available from the Zhou Lab |
| Oligo | ZZ1269 | This study | 5’-ggttGTCGACGATCCtgttgttctggaaattgag-3’ | Available from the Zhou Lab |
| Oligo | ZZ1330 | This study | 5'-aataccccggGCTATGGGCTTGTCTCCTGCCGC-3' | Available from the Zhou Lab |
| Oligo | ZZ1331 | This study | 5’-ggttaGGTACCTTATGCACAATTATTTTGGGTTGC-3’ | Available from the Zhou Lab |
| Oligo | ZZ1332 | This study | 5'-aataccccggGCTGCATTCTCCTGCAAGGATCAG-3' | Available from the Zhou Lab |
| Oligo | OC79 | This study | 5’-TTTTTTTACATGTCTCTTTTCCTTCTTCTTTCTATCCCG-3’ | Available from the Zhou Lab |
| Oligo | OC80 | This study | 5’-AAAAAAAGGATCCGCATCGCTACACCATCAGG-3’ | Available from the Zhou Lab |
